# Evolutionary crowdsourcing: alignment of fitness landscapes allows cross-species adaptation of a horizontally transferred gene

**DOI:** 10.1101/2022.09.13.507827

**Authors:** Olivia Kosterlitz, Nathan Grassi, Bailey Werner, Ryan Seamus McGee, Eva M. Top, Benjamin Kerr

## Abstract

Genes that undergo horizontal gene transfer (HGT) evolve in different genomic backgrounds as they move between hosts, in contrast to genes that evolve under strict vertical inheritance. Despite the ubiquity of HGT in microbial communities, the effects of host-switching on gene evolution have been understudied. Here, we present a novel framework to examine the consequences of host-switching on gene evolution by probing the existence and form of host-dependent mutational effects. We started exploring the effects of HGT on gene evolution by focusing on an antibiotic resistance gene (encoding a beta-lactamase) commonly found on conjugative plasmids in Enterobacteriaceae pathogens. By reconstructing the resistance landscape for a small set of mutationally connected alleles in three enteric species (*Escherichia coli, Salmonella enterica*, and *Klebsiella pneumoniae*), we uncovered that the landscape topographies were largely aligned with very low levels of host-dependent mutational effects. By simulating gene evolution with and without HGT using the species-specific empirical landscapes, we found that evolutionary outcomes were similar despite HGT. These findings suggest that the adaptive evolution of a mobile gene in one species can translate to adaptation in another species. In such a case, vehicles of cross-species HGT such as plasmids enable a distributed form of genetic evolution across a bacterial community, where species can ‘crowdsource’ adaptation from other community members. The role of evolutionary crowdsourcing on the evolution of bacteria merits further investigation.

## Introduction

In bacterial communities, genes that are transferred horizontally between species evolve in dramatically different genomic backgrounds as they move between hosts (Redondo-Salvo et al. 2020). This stands in stark contrast to genes that evolve under strict vertical inheritance, where the genomic backdrop is relatively constant over time. Even though horizontal gene transfer (HGT) is common and important in bacterial evolution, the effect of host-switching on the evolution of genes that undergo HGT (hereafter ‘mobile genes’) has received little attention.

A mobile gene that is transferred from one host species to another may exhibit changes in the fitness effects of mutations due to the new host’s genomic context. If beneficial mutations in one host are also beneficial in other hosts, then one species may effectively “crowdsource” adaptive evolution of the mobile gene. That is, a focal species that transfers a mobile gene to a different species and later reacquires that gene can benefit from adaptive genetic changes that occurred to the mobile gene while in the second host. In contrast, if beneficial mutations in one host are detrimental in other hosts, then the crowdsourcing value from HGT decreases. In this case, greater adaptive progress is expected to occur “in-house.” To determine the adaptive consequences of HGT, we need to understand how the fitness effects of mutations change in sign (beneficial or deleterious) or magnitude depending on the host harboring the mobile gene. We term this dependence *host epistasis*, a subcategory of genetic epistasis in which the fitness effects of mutations depend on the whole host genomic background rather than a subset of variable sites (Weinreich et al. 2006). If the host genome is likened to an environmental context for the evolving mobile gene, host epistasis could be envisioned as a kind of environmental epistasis (i.e., GxE interaction) (Lindsey et al. 2013). Both genetic and environmental epistasis have been shown to profoundly shape adaptive evolution (Tan et al. 2011; Goulart et al. 2013). Here, we sought to determine the presence and form of host epistasis and its effects on the evolution of a mobile gene.

A visual metaphor that can aid in understanding the evolutionary effects of host epistasis is a ‘fitness landscape’ that maps a network of mutationally connected genotypes to fitness. Host epistasis manifests as differences in the landscape topography across hosts. To illustrate different forms of host epistasis and their evolutionary consequences, we focus on a simple example involving two host species and three variant sites in a mobile gene. In Figure 1a, the landscapes of the blue and red hosts are aligned with no instances of sign host epistasis (i.e., the signs of the fitness effects of mutations are the same in the two hosts). In this scenario, HGT between the hosts does not impact the evolutionary end point reached in the red host relative to adaptation without HGT (Figure 1b). These conditions enable evolutionary crowdsourcing where the red host can make use of the transient adaptation in the blue host. In Figure 1c, the landscapes of the two hosts are mirror images of one another, indicating rampant sign host epistasis. Here, adaptation in the blue host is counterproductive to evolutionary progress in the red host (Figure 1d). This scenario highlights evolutionary “insourcing” where the red host can make more progress if adaptation remains in-house without HGT. A more subtle case is found in Figure 1e where a handful of mutations exhibit sign host epistasis. Specifically, there is a suboptimal fitness peak in the red host landscape that is absent in the blue host landscape. In this case, adaptive evolution in the blue host explores additional regions of genotype space, and HGT from the blue to red host introduces genetic variation that effectively releases the red host from a suboptimal evolutionary endpoint. This scenario highlights evolutionary “outsourcing” where HGT can qualitatively alter the evolutionary trajectory in the red host relative to adaptation without HGT (Figure 1f). These simple cases illustrate that comparing landscape topographies across hosts is the first step in determining how cross-species HGT may affect mobile gene evolution.

**Figure 1:**
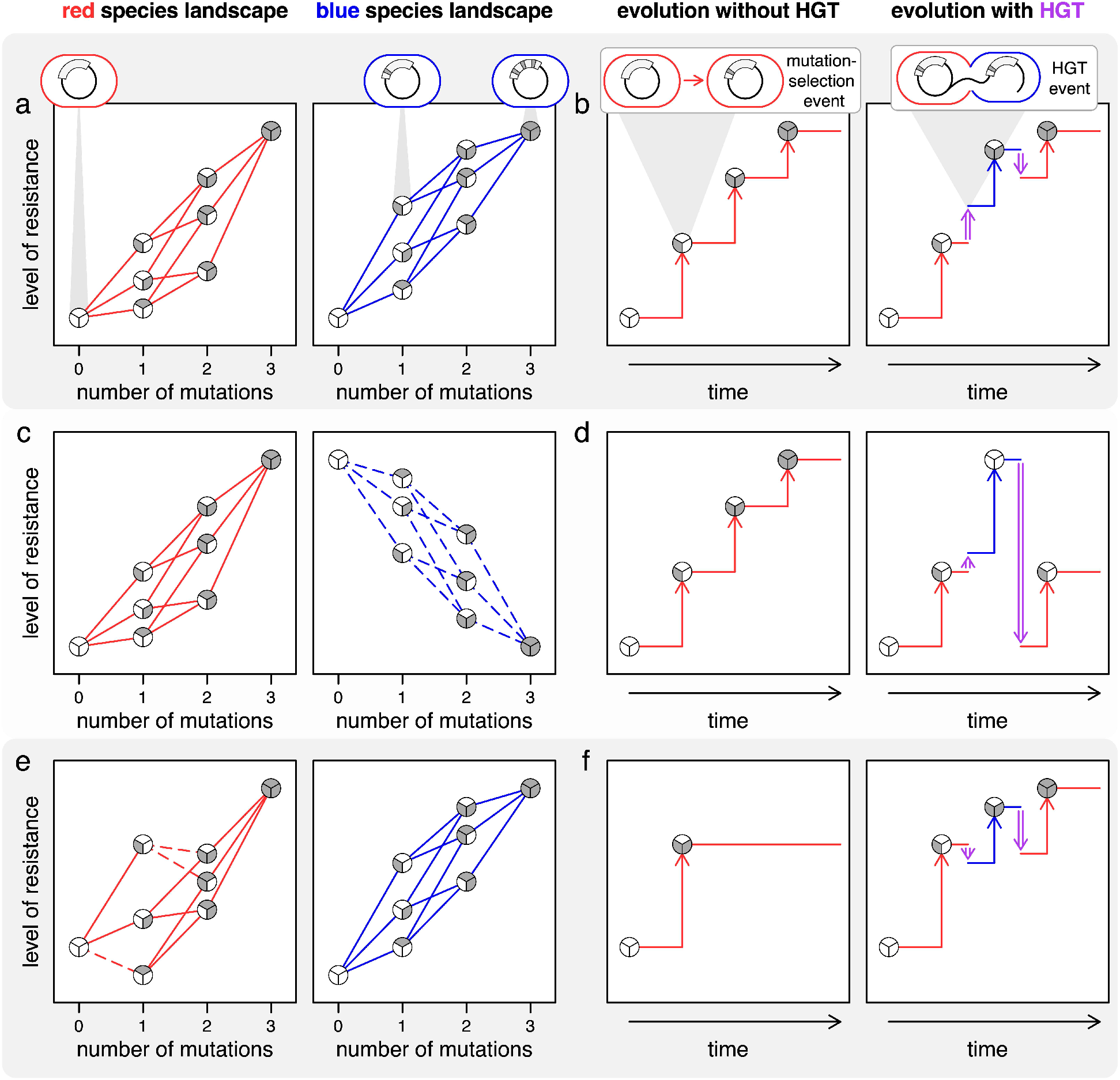
Effect of HGT on mobile gene evolution with hypothetical host-specific landscapes. Here, we consider a simple bi-allelic three-site landscape in two hosts (differentiated by the colors red and blue) for a gene encoded on a mobile genetic element. (a, c, e) The adaptive landscape can be visualized by plotting the genotype’s level of resistance (taken to be a proxy for fitness) as a function of the number of mutations on a wild-type (WT) background. Each of the 2^3^ = 8 genotypes is represented by a circle divided into ‘wedges’ equal to the number of sites (3 in this case) where the evolved variant at a site is shown by shading the wedge (in grey). The edges (lines where the color matches the host) connect genotypes differing by a single mutation. The effect of a mutation (beneficial or deleterious) is emphasized with the line type (solid or dashed, respectively). (a) In the first example, the landscapes of the red and blue host are well aligned (note there are no mutations having fitness effects of opposite sign). (b) If we assume that selection is strong and mutation is weak, we can represent the fixation of each beneficial mutation (vertical arrows) as a step up in the level of drug resistance. In this represented evolutionary trajectory after three mutational events (left panel), the population reaches the adaptive peak, a genotype from which all mutations are detrimental. When HGT (vertical purple double-ended arrow) to and from the blue host occurs preceding and following the second mutational event (right panel), the population still reaches the adaptive peak given that the landscapes (part a) of the blue and red host are aligned. This scenario is an example of evolutionary crowdsourcing where the red host can make use of the transient adaptation in the blue host. (c) Here, we show a different example where the landscapes of the two hosts possess rampant sign host epistasis. Thus, mutational steps are beneficial (solid lines) in the red host but are deleterious (dashed lines) in the blue host. (d) This scenario is an example of evolutionary insourcing where transient adaptation in the blue host is counterproductive to the evolutionary progress in the red host. (e) In this last example, the landscapes of the two hosts have only a handful of mutational steps exhibiting sign host epistasis; however, given the location of these mutations, a suboptimal fitness peak occurs in the red host landscape that is absent in the blue host landscape. (f) Evolution in the red host can result in a suboptimal evolutionary endpoint (left panel). However, adaptation in the blue host can effectively release the red host from the suboptimal endpoint (right panel), a scenario that highlights evolutionary outsourcing.

In a previous empirical study, sign host epistasis was uncovered in an antibiotic resistance landscape (Guerrero et al. 2019; Ogbunugafor and Eppstein 2019). However, this prior work focused on a small set of alleles in an essential gene primarily encoded in the bacterial chromosome. Surprisingly there has been, to our knowledge, no attention given to *mobile* genes where host epistasis may be most relevant due to HGT facilitating host-switching. Here, we experimentally constructed a portion of a mobile gene’s landscape in three Enterobacteriaceae pathogens. This gene naturally resides on conjugative plasmids in enteric bacteria and encodes a beta-lactamase in which a handful of mutations are known to increase resistance to various beta-lactam antibiotics by orders of magnitude (Weinreich et al. 2006). Our study aims to reveal the presence and nature of host epistasis and its consequences on the evolution of drug resistance.

## Experimental Approach

We assessed the host-specific landscape topography of a set of plasmid-borne antibiotic resistance alleles using a high-throughput multiplexed assay. Each allele was mapped to the level of resistance it conferred in the focal host, which is a proxy for the fitness. Together, these mappings formed a set of mutationally connected alleles comprising the “resistance” landscape. Our basic approach to assess the resistance level of each allele in each host consisted of three steps. First, each allele in the set was engineered, tagged with unique barcodes, ligated into a plasmid, and transformed into a given host (Figure 2a). Second, all transformants were pooled to create the initial host library and subsequently inoculated and incubated in a series of tubes with an increasing concentration of the relevant antibiotic (Figure 2b). Third, the pre- and post-selection cell count estimates of each allele were used to approximate the growth rate corresponding to each allele within each antibiotic concentration (Figure 2c). The estimated growth rates for each allele across the gradient yielded a dose response curve by fitting a log-logistic function. The inflection point of this curve, namely where growth rate is dropping most precipitously, was our measure of its level of resistance (similar to the minimum inhibitory concentration). Collectively, the resistance levels for the set of alleles determined the topography of the landscape for the given host (Figure 2d). By implementing this procedure (see Supporting Material for additional details) in multiple bacterial species, we were able to compare landscapes across different hosts.

**Figure 2:**
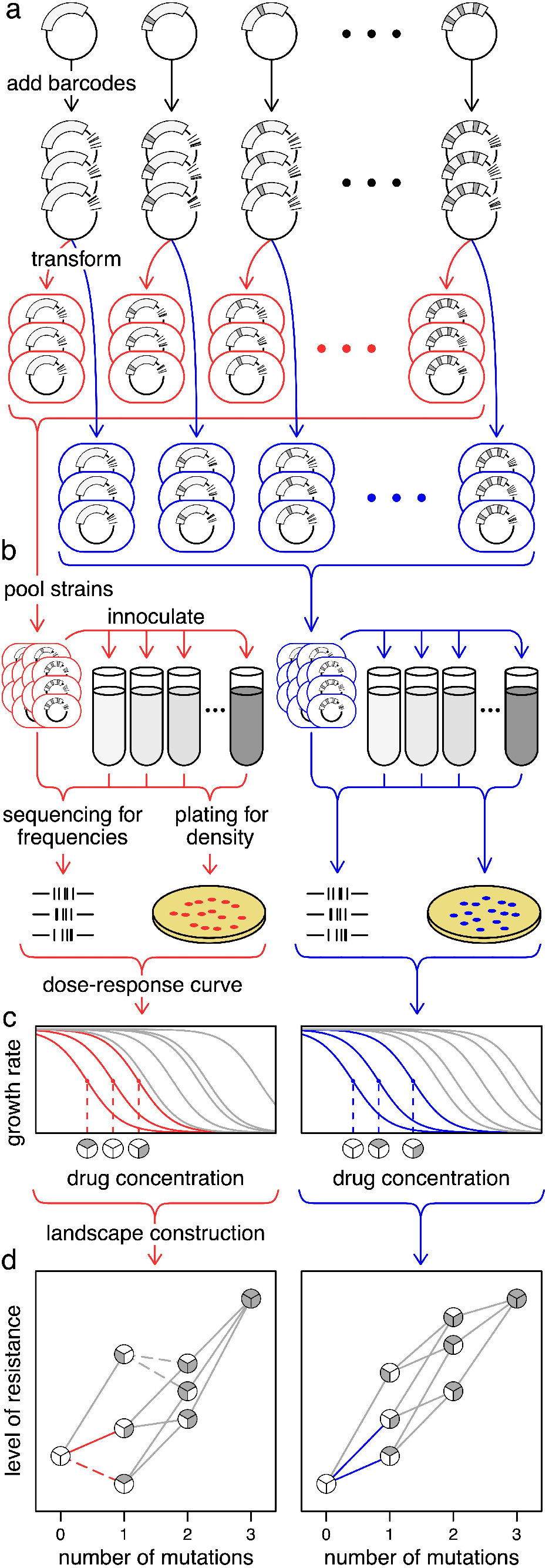
A multiplexed protocol for constructing host-specific landscapes. (a) To construct each allele of interest, mutations (dark grey notches) are introduced into a focal gene (rectangular arc) on a plasmid. To facilitate allele tracking in the experiment, each allele is tagged with three unique barcodes (black notches). Subsequently, the barcoded alleles are transformed into each host species (‘red’ and ‘blue’ hosts are shown here). (b) To assess the resistance level of each allele, all transformants within a species are pooled to create the initial bacterial library and inoculated into an antibiotic gradient (the intensity of grey-shaded medium increases with antibiotic concentration). Samples are acquired before and after incubation to determine barcode frequency using deep sequencing and the total population density using dilution plating. (c) Using the product of total population density and barcode frequencies associated with each allele before and after selection at a given concentration, a growth rate can be calculated (specific to the allele and the drug concentration). For each allele, the estimated growth rates across the antibiotic gradient yields a dose response curve by fitting a log-logistic function where the level of resistance is given by the inflection point of the curve (three alleles are highlighted per host where the resistance of each is given by the dashed vertical line). (d) The landscape topography for each host is given by the collection of the set of alleles’ resistance levels (the x-axis values for inflection points in part c). The connections between the three highlighted genotypes from part c are shown in the host-specific color.

## Results and Discussion

We constructed the resistance landscape for the *bla* gene, which encodes a beta-lactamase, in three enteric species (*Escherichia coli, Salmonella enterica*, and *Klebsiella pneumoniae*). The allele set of the *bla* gene is comprised of all combinations of five particular mutations (2^5^ = 32 alleles, nodes in Figure 3a), which in combination increased resistance in *E. coli* to cefotaxime (a beta-lactam antibiotic) by five orders of magnitude. Weinreich and colleagues previously used this allele set to construct the resistance landscape in *E. coli* (Weinreich et al. 2006). In this study, we used this allele set to determine the degree of topographic alignment across three enteric species by analyzing the existence and form of host epistasis for each mutation connecting two alleles in our set.

**Figure 3:**
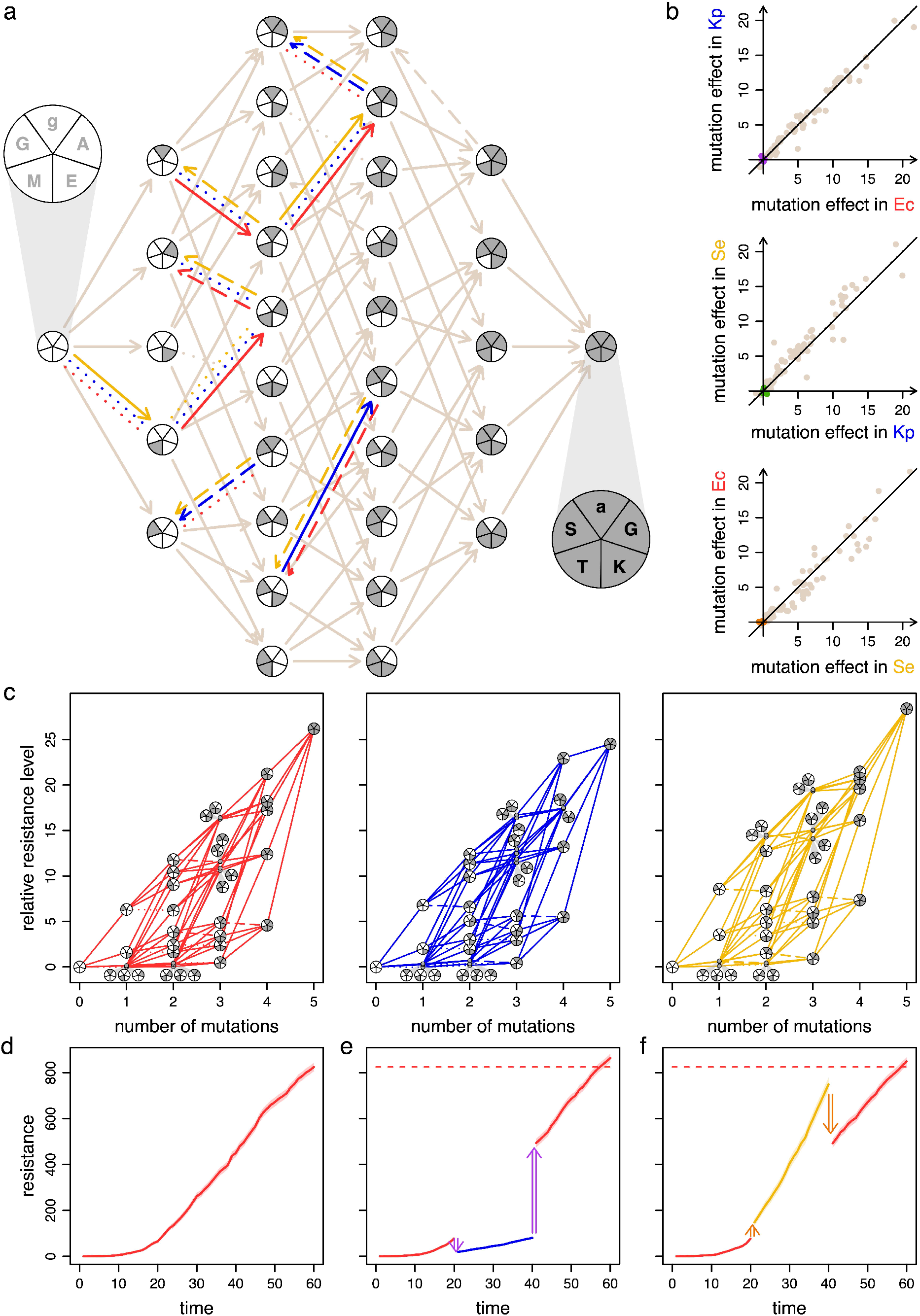
Multi-host landscapes of a mobile gene reveal evolutionary crowdsourcing. (a) The resistance landscape with five mutations to the TEM-1 allele encoding a beta-lactamase was constructed for three enteric species: *Escherichia coli* (red), *Klebsiella pneumoniae* (blue), and *Salmonella enterica* (yellow). The five mutations are displayed with five wedges in each node where shading indicates the presence of a mutation. In the pie blow ups, each mutation is indicated with either a lower- or uppercase letter corresponding to either a single nucleotide polymorphism in the promotor region or an amino acid substitution, respectively. The circle in which all the wedges are white represents the TEM-1 allele (possessing low cefotaxime resistance), whereas the circle in which all the wedges are grey represents the most resistant allele (in all species). Starting at the 12 o’clock position and moving clockwise on the pie charts, the mutations were g4205a, A42G, E104K, M182T, and G238S. The mutational steps that exhibited sign host epistasis are split into three arrows (one for each host) where the effect in each host (red, blue, and yellow edges) is indicated as beneficial (solid line), neutral (dotted line), or deleterious (dashed line). If the mutational step had the same effect in all three hosts (i.e., no sign host epistasis), then only one edge (brown) with the corresponding effect (solid, dotted, or dashed) is shown. (b) The effect of each mutational step (80 edges in the directed network in part a) on the level of resistance (akin to the slope in part c) are compared across each species pairing. The relative resistance level, 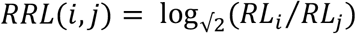, involves comparing the resistance level (*RL* in μg ml^−1^) of a focal genotype *i* to a different genotype *j*. The points plotted here give the effects of mutations in the relevant species as genotypes *i* and *j* differ by a single mutation. The mutational steps that exhibited sign host epistasis (split arrows in part a) had small effects (purple, green, orange dots near zero in the top, middle, and bottom panel, respectively) compared to the mutational steps exhibiting no host sign epistasis (brown dots). (c) Given the low number of mutational steps that exhibited sign host epistasis and their small effects, the *E. coli, K. pneumoniae*, and *S. enterica* (red, blue, and yellow, respectively) landscapes were largely aligned. Here, the *RRL(i,j*) for each host is computed by comparing the resistance level (*RL*) of a focal genotype *i* to the allele with no mutations (i.e., *j* is the TEM-1 allele). (d) Using an evolutionary simulation (see Materials and Methods for details), the average level of resistance (over 1,000 replicates with the standard error given by the shading) increased due to gene evolution in *E. coli*. (e, f) Under the same evolutionary period as part d, a similar evolutionary endpoint is reached in *E. coli* when the gene evolved in a different species during a middle period facilitated by HGT events (double-ended arrows). The dashed red line indicates the average evolutionary endpoint from part d.

Only 8 out of the 80 possible mutations exhibited sign host epistasis (red, blue, and yellow edges connecting distinct nodes in Figure 3a). Furthermore, these 8 mutations changed resistance by only small amounts (purple, green, and orange points near zero in Figure 3b), whereas all larger effect mutations exhibited similar increases in resistance across species (brown points in Figure 3b). Thus, we found that the host-specific landscapes were generally aligned (apparent in the structural similarity among the diagrams in Figure 3c).

As illustrated in Figure 1a and b, topographical congruence potentially translates to crowdsourcing of the adaptive evolution of a mobile gene. However, the landscape alignment in our three species was not perfect. As seen in Figure 1c, even a handful of mutations exhibiting sign host epistasis can impact the evolutionary trajectory of a mobile gene. To gauge the consequences of the topographical differences we found, we simulated evolution on our empirically determined landscapes (see Supporting Material for detailed description). Each simulation consisted of several rounds of stochastic mutation (limited to alleles in our set) and selection for maximal resistance (as if all alleles present were placed on a drug gradient and the allele growing at the highest concentration was picked). As a baseline, we tracked the average level of resistance (over 1,000 replicates) for genetic evolution within a single host without transfer to another species (Figure 3d, SI Figure 1a and d). To determine the effect of host-switching due to HGT, we simulated evolution over three distinct periods where several rounds of stochastic mutation and selection occurred in different species (Figure 3b-c, SI Figure 1b-c, and SI Figure 1e-f). Specifically, the mobile gene evolved in a single species (hereafter the ‘focal’ host) for several rounds in an initial period before an HGT event moved the gene into a different species (hereafter the ‘transient’ host) initializing a middle period. A second HGT event returned the gene to the focal host from the transient host initializing a third and final period. Despite time evolving in a transient species, the final level of resistance for the same total period of evolution was statistically indistinguishable (SI Figure 2). This pattern was maintained when HGT occurred between different species at different periods with various simulation parameters (SI Figure 6 SI Figure **8**). Thus, our species can effectively crowdsource the evolution of antibiotic resistance in this case.

For both the landscape reconstruction and evolutionary simulation, there are a few caveats that warrant attention. First, we focused on a small set of mutations in one gene in three closely related species. It is possible that more or less cross-species topographical alignment could occur when considering additional or alternative mutations in the *bla* gene, a different gene, or different species (e.g., hosts that are more phylogenetically distant). Second, our “fitness” landscapes mapped each allele to its level of resistance, but the fitness of any genotype will be affected by more than resistance (Ogbunugafor et al. 2016). For instance, genotypes may exhibit different baseline growth rates (i.e., in drug-free conditions), which may not correlate with drug resistance (Das et al. 2020). Third, our evolutionary simulation makes several simplifying assumptions. Specifically, we assumed a series of selective sweeps (where a more resistant mutant replaces its immediate ancestor) punctuated by cross-species HGT. However, multiple genotypes within and across species can compete in natural bacterial communities, and the outcomes of these competitions (as well as the opportunities for HGT) will be governed by the distribution of the relevant species across a potentially heterogeneous environment (e.g., a multi-species biofilm in a drug gradient). In addition, our simulations glossed over some of the distinguishing features of conjugative plasmids, such as multiple copies per cell, fitness costs, and basic rates of conjugation and segregational loss. Many of these plasmid features vary with host context (De Gelder et al. 2007; Dimitriu et al. 2019; Kosterlitz et al. 2022) and these differences may influence HGT opportunities and competitive outcomes. Considering these limitations, more complex simulation frameworks incorporating factors such as growth differences, clonal interference, environmental heterogeneity, and basic life history traits of plasmids would be promising directions for future work.

Despite these noted caveats, evolutionary adaptation of our focal mobile gene can be a cosmopolitan affair, in which the progress made in one species readily translates to progress in another. We emphasize that the availability of widespread evolutionary crowdsourcing in a microbial community through HGT will be affected by the prevalence of host sign epistasis, its effect size, and its topographical location. Indeed, we subjected the multi-host landscapes of an essential chromosomal gene (SI Figure 3) from a recently published study (Guerrero et al. 2019) to our evolutionary simulation framework showing that occurrence of HGT in this case would have hindering (as in Figure 1b) or facilitating (Figure 1c) effects depending on the focal and transient host (SI Figure 4SI **Figure 5**). We find it interesting that the level of sign host epistasis and the accompanying effect sizes for this essential gene are greater than the level for our nonessential gene, which is commonly found on plasmids. This pattern highlights a connection to the “complexity hypothesis,” in which the proteins encoded by genes experiencing higher rates of HGT are less connected to other proteins in the cell (Jain et al. 1999; Novick and Doolittle 2020). If fewer intracellular connections translate to fewer opportunities for host-dependencies, it would not be surprising if these more “modular” mobile genes also exhibit less sign host epistasis. To gauge whether evolutionary crowdsourcing is more often an option for genes that experience a high rate of HGT, it will be necessary to construct landscapes for additional genes undergoing different rates of HGT across the same set of species (i.e., controlling for phylogenetic relatedness).

In summary, we present a novel framework to examine the molecular evolution of mobile genes – a highly relevant subset of genes evolving with an additional mode of genetic transmission (i.e., horizontal). In this study, we uncovered that for a small set of mutations in a common mobile gene, landscape topography and thus evolutionary outcomes are largely aligned across closely related species. These findings suggest that mobile genes adapting in one species can lead to adaptation for another species. In such a case, conjugative plasmids and other vehicles of cross-species HGT will enable a distributed form of genetic evolution across a bacterial community, where any particular species can crowdsource adaptation from other community members.

## Methods, Data, and Code Availability

Detailed Materials and Methods are available in the Supporting Material.

## Acknowledgements

This work was supported by the National Institute of Allergy and Infectious Diseases Extramural Activities from the National Institutes of Health and Division (grant number R01 AI084918 to E.M.T. and B.K.), the Environmental Biology Division from the National Science Foundation (grant number 2142718 to B.K. and E.M.T.), and a Graduate Research Fellowship from the National Science Foundation (grant number DGE-1762114 to O.K.).

## Supporting Material

### Materials and Methods

#### General reagents

Unless otherwise noted, all enzymes and related buffers were obtained from New England Biolabs. Plasmid isolation kits were obtained from Qiagen. DNA cleaning and gel extraction kits were obtained from Zymo Research. Oligonucleotide primers were obtained from Integrated DNA Technologies. Sanger sequencing was conducted by GENEWIZ from Azenta Life Sciences.

#### Allele construction and barcoding

To generate the beta-lactamase alleles, we mutated the cloning plasmid, pBR322, using the New England Biolabs Site-Directed Mutagenesis Kit per manufacturer’s instructions. The plasmid contains two accessory genes, the focal *bla* gene and a tetracycline resistance gene, *tetA*. Plasmid maintenance was ensured by supplementing the culture medium with 15 μg ml^−1^ tetracycline. TEM-1 was the starting allele for the *bla* gene and used as the template for mutagenesis. The first round of cloning created the five single variant alleles. For each variant, a custom pair of primers was designed with the mutation coded in the forward primer. All primers are listed in SI Table 1. Each engineered variant was isolated and variant sequences were confirmed with Sanger sequencing. All Sanger sequencing primers are listed in SI Table 2. The double, triple, quadruple, and quintuple variant alleles were constructed using the same procedure. All engineered alleles are listed in SI Table 3.

Each beta-lactamase allele was associated with three unique molecular barcodes. To facilitate the engineering of barcodes, we modified the backbone of the pBR322 plasmid before generating the beta-lactamase alleles (described above) by adding two restriction sites, Nsil and NcoI, downstream of the *bla* gene. To barcode each allele, we digested the mutated plasmids with Nsil and NcoI at 37°C for 1 h and the restriction enzymes were heat inactivated at 65°C for 20 min. We isolated the digested vector backbone using a gel extraction kit and purified the DNA. We next prepared the double-stranded barcoded fragments to be inserted by ligation using two oligonucleotides: (1) an oligonucleotide with 18bp random barcode sequences nested between the Nsil and NcoI cut sites to be used in directional cloning, and (2) a shorter priming oligonucleotide containing homology to the barcode oligonucleotide. These two oligonucleotides are listed in SI Table 4. To construct the double-stranded barcode fragment, we mixed 1 μL of each oligonucleotide with 5 μL of CutSmart Buffer and 47.5 μL of ddH2O and annealed these oligonucleotides together by incubating at 98°C for 3 min followed by a ramping down to 25°C at −0.1°C/s. After annealing, we added 1 μL of Klenow polymerase (exonuclease negative) and 1.65 μL of 1mM dNTPs to make the barcode fragment double stranded by incubating at 25°C for 15 min, 75°C for 20 min, and then a ramping down to 37°C at −0.1°C/s. We digested the double-stranded fragment using the same enzymes and protocol for digesting the vector backbone described above. The digested barcoded fragment was purified. The digested vector and barcoded fragment were ligated at 21°C for 30 min, the enzymes were heat inactivated at 65°C for 10 min and the circular products were transformed into *E. coli*. To get three barcodes per beta-lactamase allele, we isolated and sequenced the barcode sequences in three colonies with Sanger sequencing. Given the 32 alleles with 3 barcodes per allele, the beta-lactamase library contained 96 engineered plasmids.

To create the heterogenous populations for each host species, we first transformed each of the 96 engineered plasmids independently into each host, isolated a colony from each transformation, and stored at −80°C in 15% (v/v) glycerol. For each species, the 96 strains were grown overnight, mixed at equal volumes, and stored at −80°C in 1 ml aliquots to create library stocks for each species.

#### Bacterial Strains, Media, and Culture Conditions

Hosts included three *Enterobacteriaceae* species: *Escherichia coli* DH10B (Durfee et al. 2008), *Klebsiella pneumoniae* Kp08 (Jordt et al. 2020), and *Salmonella enterica* serovar *typhirium* LT2 (McClelland et al. 2001). We use the first letter of the genus and species name (Ec, Kp, and Se, respectively) to refer to these species throughout. All strains were cultured at 37°C in lysogeny broth (LB).

#### Pooled competitions assays

The resistance level conferred by the *bla* alleles was estimated using a modified version of a standard minimum inhibitory concentration assay (Wiegand et al. 2008). Briefly, a 1 ml library stock was thawed and added to 50 ml of growth medium supplement with 15 μg ml^−1^ tetracycline and grown overnight. The library was diluted to an initial cell density close to 10^5^ cell ml^-1^. We plated the diluted libraries in triplicate to assess the actual initial cell density which is reported in SI Table 5. To start competitions assays, 2.5 ml of diluted library was inoculated into 41 test tubes with 2.5 ml medium. One test tube contained growth medium without CTX. The other 40 test tubes were supplemented with CTX according to a drug-gradient from 0.00393 up to 2049.37 μg ml^-1^ using √2-fold dilutions. After overnight growth in the 41 test tubes, 1ml samples were taken from the test tubes with visible turbidity for library amplification and sequencing. These samples were harvested by centrifugation and the cell pellets were stored at −20°C. Final density was also determined for each of the test tubes with visible turbidity reported in SI Table 6 using dilution plating.

#### Library amplification and sequencing

Plasmids were extracted from the cell pellets stored at −20°C and the barcode region was amplified using the primers homologous to the plasmid backbone with the following conditions: 95°C for 3 min, five cycles of 98°C for 15 s, 65°C for 15 s, 72°C for 30 s, and 72°C for 1 min. Amplicons were then purified with AMPure XP beads (Beckman Coulter) at 1:1 ratio. Each sample’s purified PCR product was amplified with a unique pair of forward and reverse indexing primers plus SyberGreen with the following PCR conditions on a miniOpticon (Bio-Rad): starting with 95°C for 3 min, fifteen cycles of 98°C for 20 s, 60°C for 15 s, 72°C for 30 s, and finishing with 72°C for 2 min. Using the relative fluorescence units, the amplicons were mixed, gel extracted, quantified by Qubit fluorometry, and sequenced on the Illumina NextSeq500 platform by the Microbial Genome Sequencing Center using custom sequencing primers. All PCR and custom Illumina sequencing primers are listed in SI Table 7.

#### Library sequence analysis

To determine barcode frequency, the 18bp barcode from each sequenced read was extracted from the FASTQ file and clustered into groups using Bartender (Zhao et al. 2018). The barcode clusters were matched to the barcodes used in this study (listed in SI Table 8). Barcode frequency was calculated by dividing the barcode counts by the total counts in the sample. Next, the approximate growth rate for each barcode was calculated using the estimated initial and final cell densities (SI Table 5 and SI **Table 6**) along with initial and final barcode frequencies using equation [1.1] (see SI Section 1 for details). Given that we have three barcodes per allele, these served as internal replicates in the competitions. We eliminated the most deviate barcode for each allele by calculating the pair-wise differences between the barcode approximate growth rates, summed the differences across the concentrations, and found the most globally deviant barcode. For each of the remaining two barcodes per allele, we fit a four-parameter log logistic dose-response curve using the drc package in R (Ritz et al. 2015). Given that there was an inflation of the bottom asymptote (i.e., non-zero), we took the lower asymptote parameter average for each species using approximately half of the alleles with the lowest MIC (n = 17). For each barcode, we used these species-specific lower asymptote values to fit a three-parameter log logistic dose-response curve. By fixing the lower asymptote, it established the same “no-growth” baseline across the alleles which improved the curve-fitting. We used the inflection point, namely where growth rate is dropping most precipitously, of the dose-response curve as the proxy for the level of resistance for each allele. In the event the lower asymptote was zero, then our resistance level would be an IC50 value. Since all the barcodes were pooled and thus exposed to the same environments, using the relative inflection points as the proxy for level of resistance was justified.

For the less resistant alleles, the concentration gradient was truncated to the highest concentration where the approximate growth rate of the most resistant allele was unaffected by the antibiotic. For the most resistant allele (g4205a, A42G, E104K, M182T, G238S) and the second most resistant allele (g4205a, A42G, M182T, G238S), the entire concentration gradient was used for fitting the three-parameter log logistic dose-response curve to obtain the inflection point for these alleles. The three-parameter estimates for the dose-response curve (upper asymptote, steepness, and inflection point) for each barcode-allele-species combination are given in SI Table 1.

To construct the host-specific landscapes (Figure 3), the level of resistance for each neighboring allele (two estimates per allele given in SI Table 1) were compared to determine if a mutational step was beneficial, deleterious, or neutral (solid, dashed, or dotted lines in Figure 3a and c). If the resistance estimates for the single mutant neighbor were all higher than the focal genotype, the mutational step was beneficial. If the estimates for the single mutant neighbor were all lower than the focal genotype, the mutational step was deleterious. If the estimates for the single mutant neighbor overlapped with the focal genotype, the mutational step was neutral. Each mutational step was compared across the three species to calculate the amount of sign host epistasis. A mutational step exhibited sign host epistasis if the effect (beneficial, deleterious, or neutral) of the mutation was different in one of the three species.

#### Evolutionary Simulations

Each evolutionary simulation comprised of periods of host-specific evolution, in which the focal gene evolved for a specified number of time steps inside a single host species (SI Table 10). An HGT event was defined as a switch in the host of the evolving gene, which occurred at specified time steps (SI Table 10). Therefore, a simulation for a gene evolving in different hosts over time consisted of distinct periods of single-species evolution linked together by HGT events.

Within each period, mutation and selection occurred at each time step. Specifically, we considered a set of random single mutants along with the ancestral allele where the genotype conferring the highest resistance fixed. We note that at some time steps the most resistant genotype may have been the ancestral allele if no mutants were generated (which becomes more likely as the mutation rate decreases) or if no generated mutant was more resistant than the ancestor (which becomes more likely as fewer mutational neighbors improve resistance). Technically, we simulated this mutation-selection process by computing the probability of genotype *j* fixing given that genotype *i* is the current ancestor. This set of probabilities depended on mutation rate, population size, and the fitness’s of genotypes *í* and *j* (see 0 for details on calculating the probabilities). If a mutation exhibited sign host epistasis, or a mutation exhibited magnitude host epistasis shifting the rank ordering (based on resistance) of the mutational neighbors, the evolutionary trajectory of a gene could be affected by HGT. Thus, sign or magnitude host epistasis could enable HGT to impact the path and endpoint of adaptive evolution.

We note that our simulation approach treats the evolutionary trajectory of a gene as a sequence of single allele states, where probabilistically the most beneficial allele of those mutationally available fixes after each time step. That is, we did not model the process of one allele displacing another over multiple time steps. Therefore, differences in allelic replacement rates across species (the time for a selective sweep) or the potential for multiple alleles to coexist (forms of clonal interference) were not incorporated into this approach. One way to think of our simulation is to imagine a population distributed across a drug gradient and the most resistant genotype is selected at every discrete time step to initiate the next round of population growth and placement along the drug gradient, which is a reasonably approximation of some directed evolution schemes (Salverda et al. 2010; Packer and Liu 2015; Salverda et al. 2017).

The empirically determined host-specific landscapes provided the information about the beneficial mutants available at each focal genotype (number and ranking). Since each genotype for the *bla* gene had multiple (replicate) estimates for its level of resistance (SI Table 9), at each time step, the resistance for each allele was sampled randomly from the set of estimates. Therefore, for each time step of the simulation, the host landscape could potentially shift; however, these shifts were small given that the variance across the resistance estimates in our assay were generally very low.

To determine the effects of various simulation parameters on the evolutionary patterns drawn for the *bla* gene (Figure 3 and SI Figure 1–SI **Figure 2**) and DHFR gene (SI Figure 4–SI **Figure 5**) from the baseline simulations, we expanded the evolutionary simulations and ran sets of simulations sweeping through parameter values. Each simulation examined a parameter in isolation by manipulating the relevant parameter (mutation rate, cumulative time, and number of simulation replicates). For the “baseline” simulations, we used a “baseline” set of parameters (mutation rate of 5 × 10^-5^, populations size was 1000, and the number of replicate simulations was 1000). Given that the *bla* and DHFR gene differed in the number of mutations (5 and 3, respectively), the cumulative time for the baseline parameters was gene-specific (60 and 36, respectively) where the amount of time steps per mutation was proportional between the two genes. For treatments with HGT, the middle period was always one third of the cumulative time. Significance was determined using a Wilcox test with a Bonferroni correction. The set of simulations and parameters used are given in SI Table 10. We demonstrate that the trends shown for the *bla* gene for the baseline condition are also observed with parameter sweeps through mutation rate (SI Figure 6), cumulative time (SI Figure 7), and number of simulation replicates (SI Figure 8). In addition, the trends shown for the DHFR gene with the baseline condition largely holds with parameter sweeps through mutation rate (SI Figure 9), cumulative time (SI Figure 10), and number of simulation replicates (SI Figure 11).

### SI Section 1: Derivation of the approximate growth rate

Here, we describe the growth rate metric used in this study. A standard growth rate equation was problematic given that the assay was multiplexed where all the alleles in our set were growing together in each antibiotic concentration that was tested. Since more alleles survived lower antibiotic concentrations compared to higher antibiotic concentrations, the number of alleles competing together decreased as the antibiotic concentration increased. Since the initial density per allele was relatively constant across the antibiotic gradient, as the antibiotic concentration increased the growing time per allele increased given less competitors and more access to resources. Here we derive an approximate growth rate metric to account for the variable growth period across the gradient. This approximate growth rate metric as a function of antibiotic concentration generated a standard dose-response curve for each allele as described in the materials and methods.

Let *c* * be the highest concentration where the genotype with the highest resistance does not see a drop in growth. That is, for *c* > *c\**, the most resistant genotype will drop in its estimated growth rate, indicative that the genotype was affected by the antibiotic. We label this most resistant genotype as *g′*. We let the growth rate calculated for genotype *g* at concentration *c* be given by 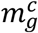. To calculate this growth rate, we used a time period of 24 hours even though the culture could have been growing for less than 24 hours. We let 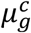 be the actual growth rate of genotype *g* at concentration *c*. Now at concentration *c*\*, we will assume that for genotype *g′*

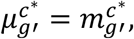

which is simply assuming that growth is occurring during the full *T* = 24 hours.

For the time being, we will assume that this growth rate remains constant for lower drug concentrations.

Now, consider some *c* < *c\**. We have the following:

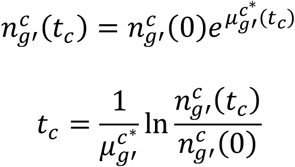

If 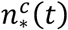 is the total cell count at time *t*, and 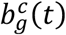 is the proportion of barcodes associated with genotype *g* in concentration *c* at time *t*, then we have

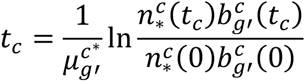

Or, assuming that the number and proportion of cells does not change from *t_c_* to *T*:

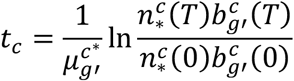

Now, we can consider any arbitrary genotype:

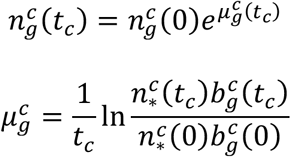

Or, assuming that the number and proportion of cells does not change from *t_c_* to *T*:

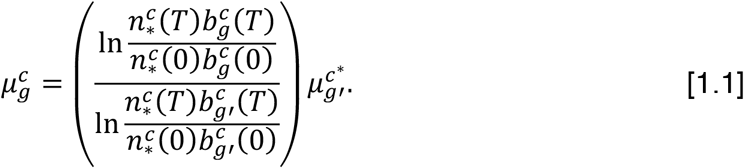

To derive equation [1.1], we assumed that growth is occurring for a full 24 hours when considering 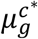 and that this growth rate is constant for lower drug concentrations. However, this second assumption is likely misplaced. Suppose for some *c* < *c\** the actual growth time is *t* *, then we have the following:

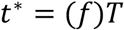

where the actual growth time is some fraction *f* of the full time period (*T* = 24 hours). Therefore,

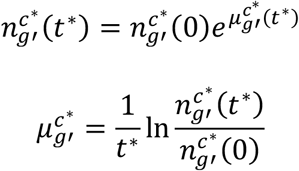

So, substituting *t*\* with (*f*)*T* where *T* = 24 and the number of cells does not change from *t*\* to *T* (the same assumption as above):

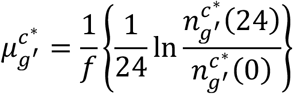

Thus, the connection between the measured growth rate 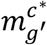 and the actual growth rate 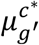, is off by a factor:

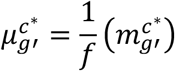

For simplicity, we assume *f* = 1 in our data analysis.

### SI Section 2: Probabilities that a given mutation is the most resistant under gradient selection

We modeled mutation and selection using a first-order Markov process. First, we describe the probability that the population shifts from the ancestral genotype (hereafter focal genotype) to a single mutant neighboring genotype at the end of a time step. We let *μ* be the probability a mutation arises in the focal genotype. The probability of mutation is small such that we ignore any instance of two or more distinct mutations and assume that most individuals within the population have the focal genotype (and the rare mutants have only a single mutation). All possible single mutant genotypes are denoted by the set **M** ≡ {*m*_1_, *m*_2_, *m*_3_, … *m_y_*} (where *y* ≡ |**M**| is the number of mutants). Focusing on one mutant genotype (e.g., *m_j_*), if the focal genotype is copied *n* time (independently), the probability that there are *k* mutants with this genotype will be:

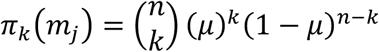

The probability that there are one or more *m_j_* individuals is then:

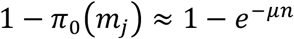

We denote this probability as *π*\* (*m_j_*).

After mutation, the population undergoes growth followed by selection across an antibiotic gradient. Here we assume that there is enough growth of the population such that if a mutant genotype is present, it is distributed across the entire antibiotic gradient. For a given focal genotype *i*, and a set of neighboring mutant genotypes labeled **M**, we denote the genotypes in set **M** that are more resistant than the focal genotype as **H_M_**(*i*). First, we start with the probability that the most resistant genotype is the focal genotype *i*. In this case, the probability is:

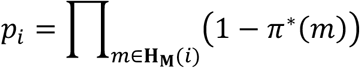

If the focal genotype is the most resistant, then this probability is defined to be one. If a mutant neighbor is more resistant than the focal genotype, this probability becomes less than one. Generally, as the focal genotype becomes less resistant, it becomes less probable for the population to stay at this genotype.

Next, we turn to the probability that the most resistant genotype comes from the mutant set **M**. Here we need to introduce more notation. For a given genotype *g*, and a set of genotypes labeled **S**, we denote the genotypes in **S** that are equally resistant to genotype *g* as **E_s_**(*g*). If genotype *g* is in the set **S**, we will define **E_s_**(*g*) to *not* include genotype *g* (i.e., *g ∉* **E_s_**(*g*)). Therefore, **E_s_**(*g*) are genotypes with equivalent resistance to genotype *g, other than g itself*. Consider a mutant genotype *m_j_*, which has higher resistance than the focal genotype *i*. We denote the set of other mutants that have higher and equivalent resistance to genotype *m_j_* as **H_M_**(*m_j_*) and **E_M_**(*m_j_*), respectively. The probability that genotype *m_j_* is chosen is given by:

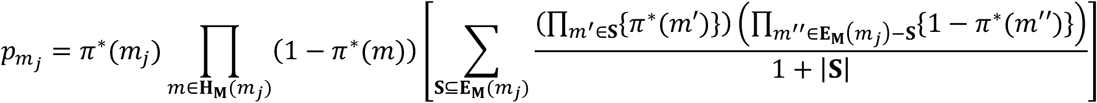

If there are never any ties between genotypes with regards to resistance, then |**E_M_**(*m_j_*)| = 0 and the above equation simplifies to:

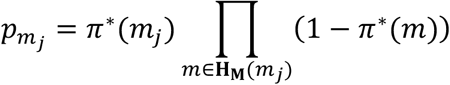

If a particular mutant *m_j_* is the most resistant mutant of the focal genotype (|**H_M_**(*m_j_*)| = 0) then the probability of picking the mutant is *π*\*(*m_j_*) which depends only on the population size and the mutation rate. More generally, the probability of selecting mutant *m_j_* covaries positively with its ranking in the set (i.e., the more resistant this mutant genotype is relative to the other mutants of the focal genotype, the more likely it is to be selected).

### Supplemental Figures and Tables

**SI Figure 1:**
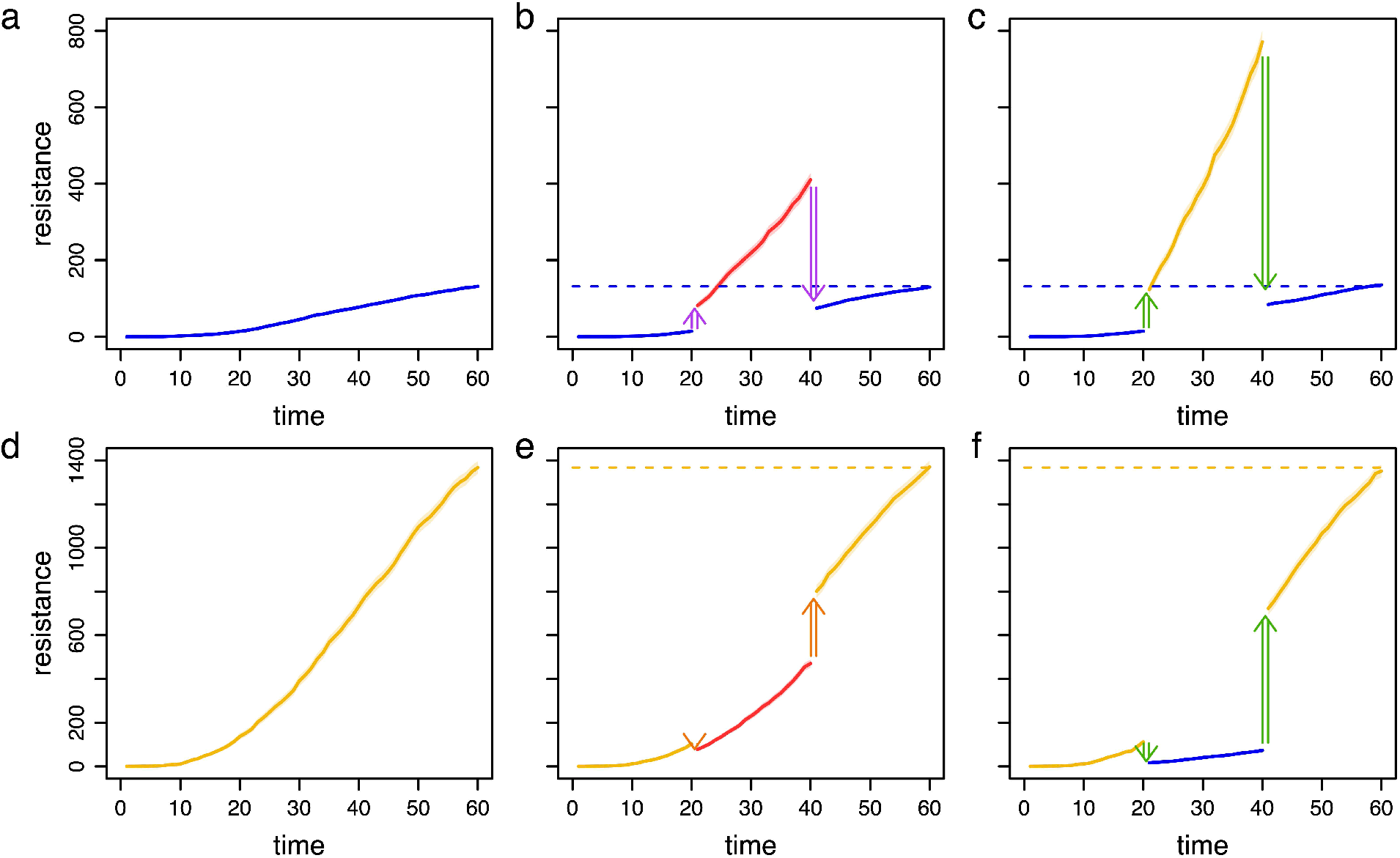
Evolutionary simulations in *K. pneumoniae* and *S. enterica* where the mobile gene evolved without (a and d, respectively) and with HGT (b, c, e, and f, respectively). The simulation framework and graphical representation is the same as Figure 3. For both *K. pneumoniae* and *S. enterica*, a similar evolutionary endpoint is reached when the gene evolved in a different species during a middle period facilitated by HGT events.

**SI Figure 2:**
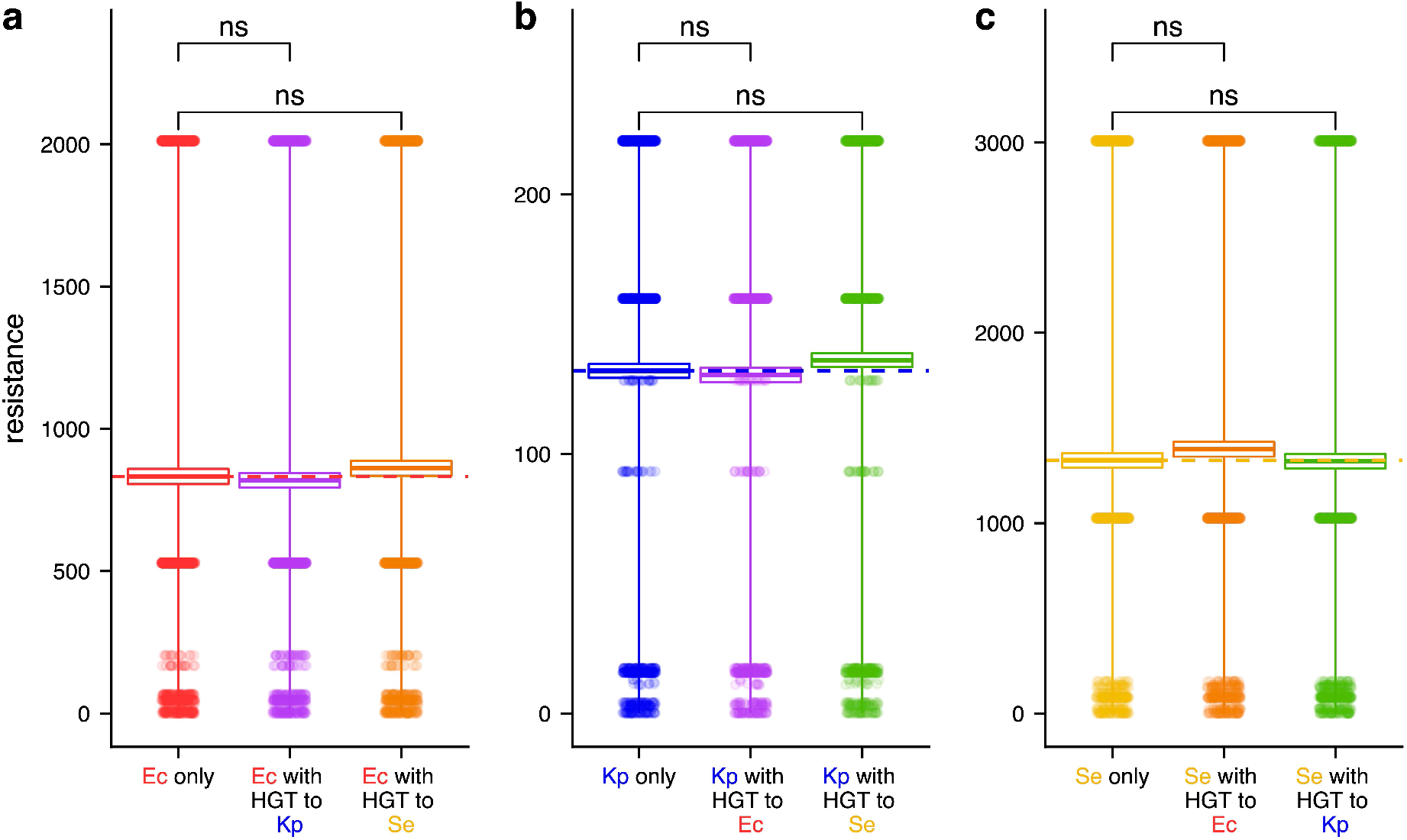
Similar evolutionary endpoints are reached when the mobile gene evolved with and without HGT. Each box summarizes the mean and standard error of the endpoint resistance values from 1000 replicates from an evolutionary simulation, which corresponds to the evolutionary endpoints in Figure 3d-f and SI Figure 1. In the baseline simulations (labelled ‘Ec only’, ‘Kp only’, and ‘Se only’), evolution of the gene occurred in the focal species (*E. coli, K. pneumoniae, and S. enterica*, respectively) for the entire evolutionary period and there is no transfer to another species. In the simulations with HGT, the mobile gene evolved in a transient species during the middle period. All comparisons to the baseline simulations were not significant by a Wilcox test with a Bonferroni correction (p=1, p=0.79, p=1, p=0.57, p=0.54, and p=1, a to c from top to bottom respectively).

**SI Figure 3:**
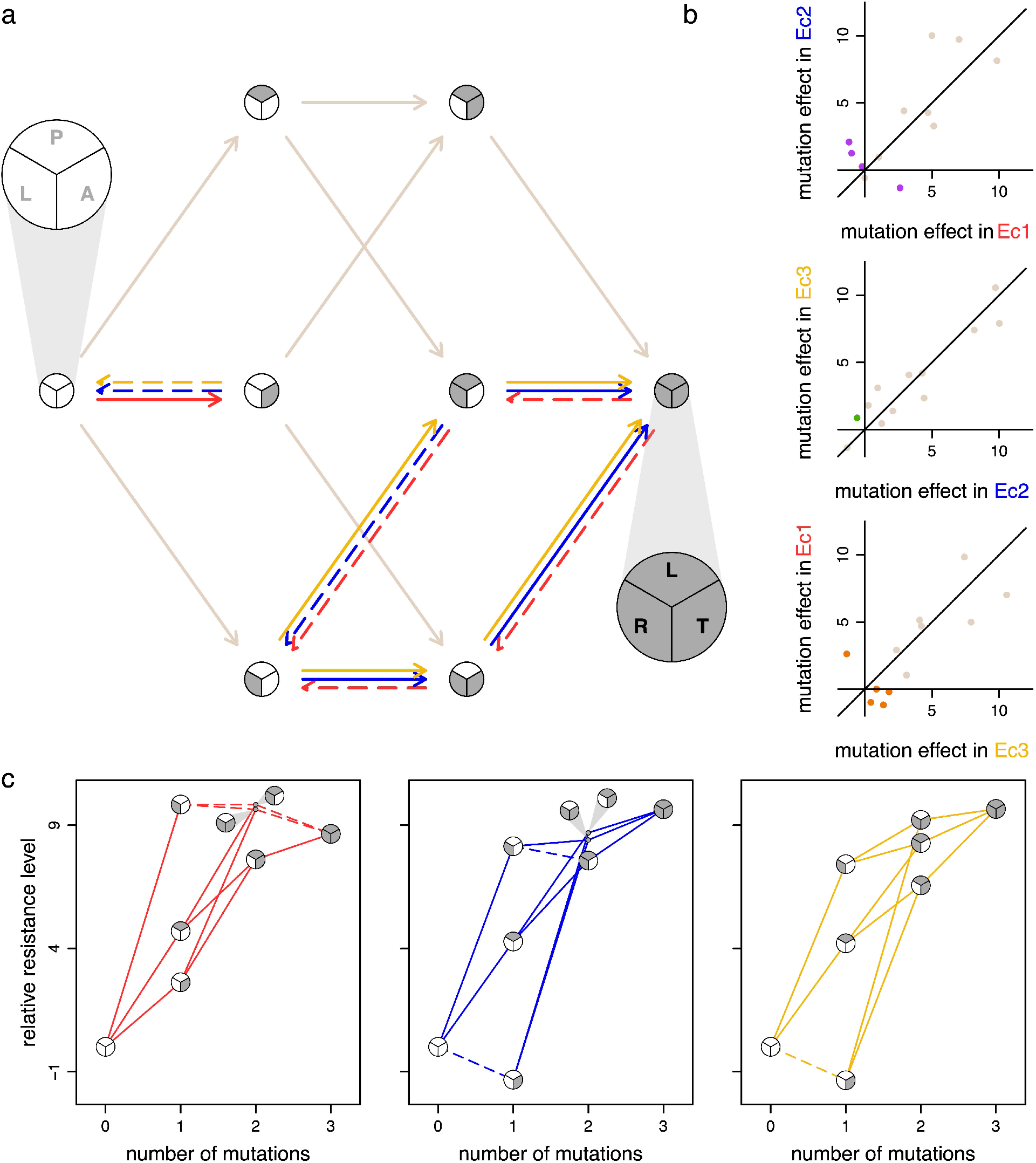
Multi-host landscapes of an essential gene from the study by Guerrero and colleagues (Guerrero et al. 2019). The graphical representations are the same as Figure 3. (a) The resistance landscape with three mutations to an *E. coli* allele of DHFR was constructed in three strains of *E. coli* referred to as ‘Ec1’, ‘Ec2’, and ‘Ec3’ (red, blue, and yellow, respectively). In the pie blow ups starting at 12 o’clock, the mutations are P21L, A26T, and L28R. (b) The effect of each mutational step (12 edges in the directed network in part a) on the level of resistance are compared across each host pairing. (c) The host-specific adaptive landscapes visualized by plotting the relative resistance level (relative to the WT background of each host) as a function of the number of mutations.

**SI Figure 4:**
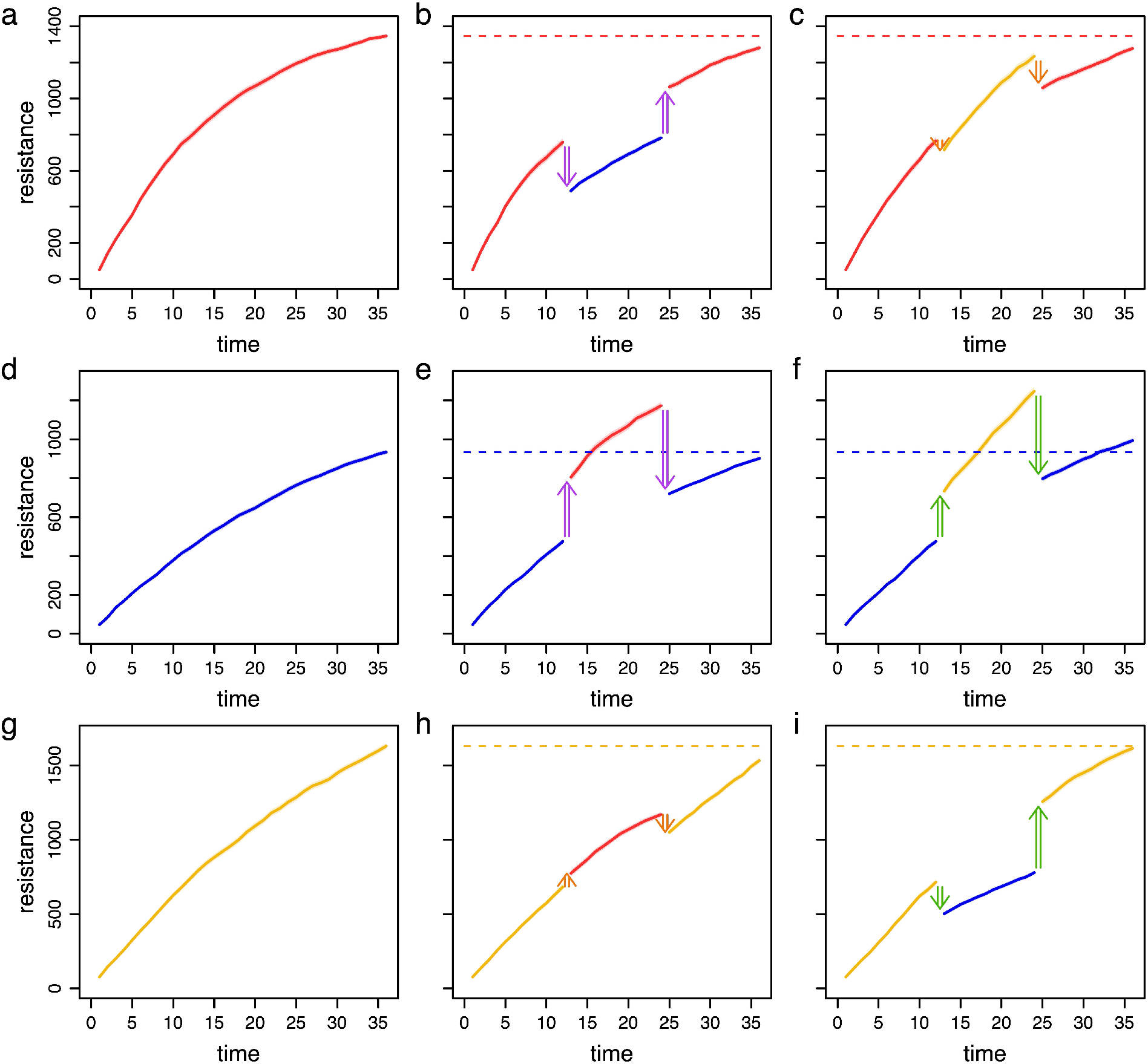
Evolutionary simulations with and without HGT using the DHFR data set from Guerrero and colleagues (Guerrero et al. 2019). The simulation framework and graphical representation is the same as Figure 3d and SI Figure 1.

**SI Figure 5:**
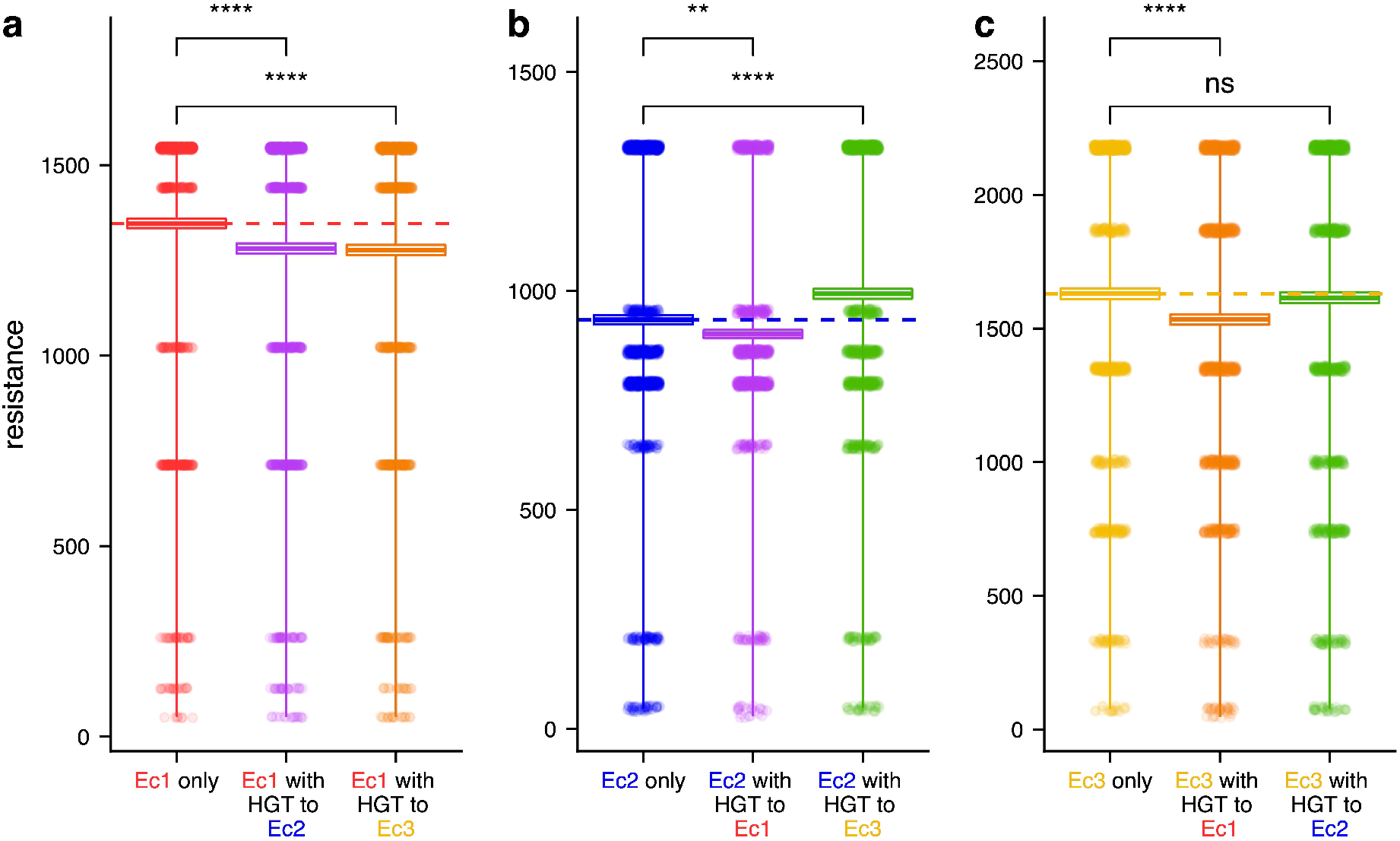
Different evolutionary endpoints are reached where the mobile gene evolved with and without HGT. The graphical representation is the same as SI Figure 2. The data corresponds to the evolutionary endpoints in SI Figure 4. The asterisks indicate statistical significance by a Wilcox text with a Bonferroni correction (two and four asterisks convey p-values in the following ranges: 0.001 < p < 0.01, and p < 0.0001, respectively), and ‘ns’ indicates statistical non-significance (p=0.71 in part c).

**SI Figure 6:**
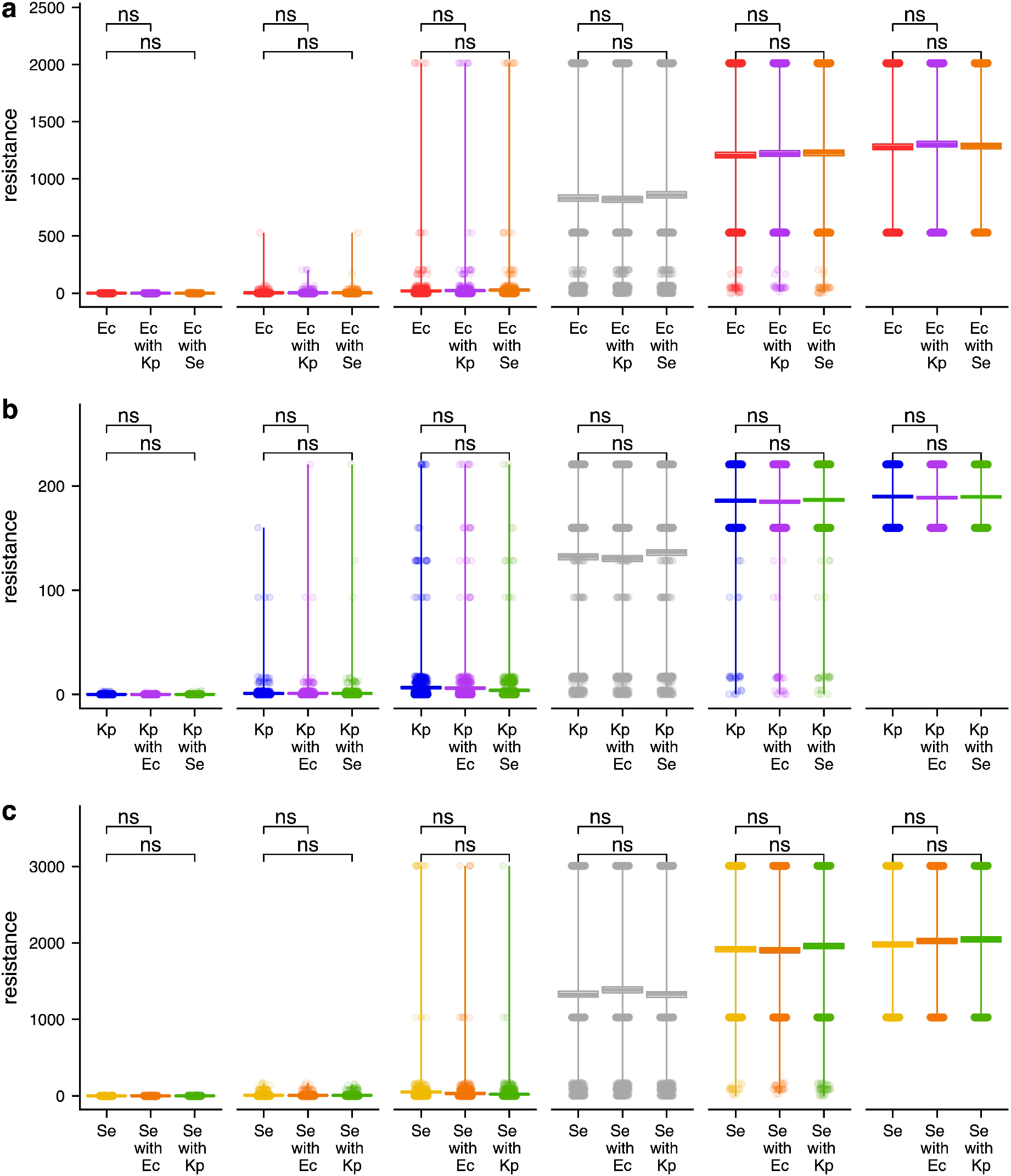
The effect of mutation rates on the evolutionary trends of the *bla* gene. The graphical representation is the same as SI Figure 2. The mutation rate increases from left to right (1 × 10^6^, 5 × 10^6^, 1 × 10^5^, 5 × 10^5^, 1 × 10^4^, and 5 × 10^4^). The gray triplicate in each part indicates the baseline simulation given in SI Figure 2.

**SI Figure 7:**
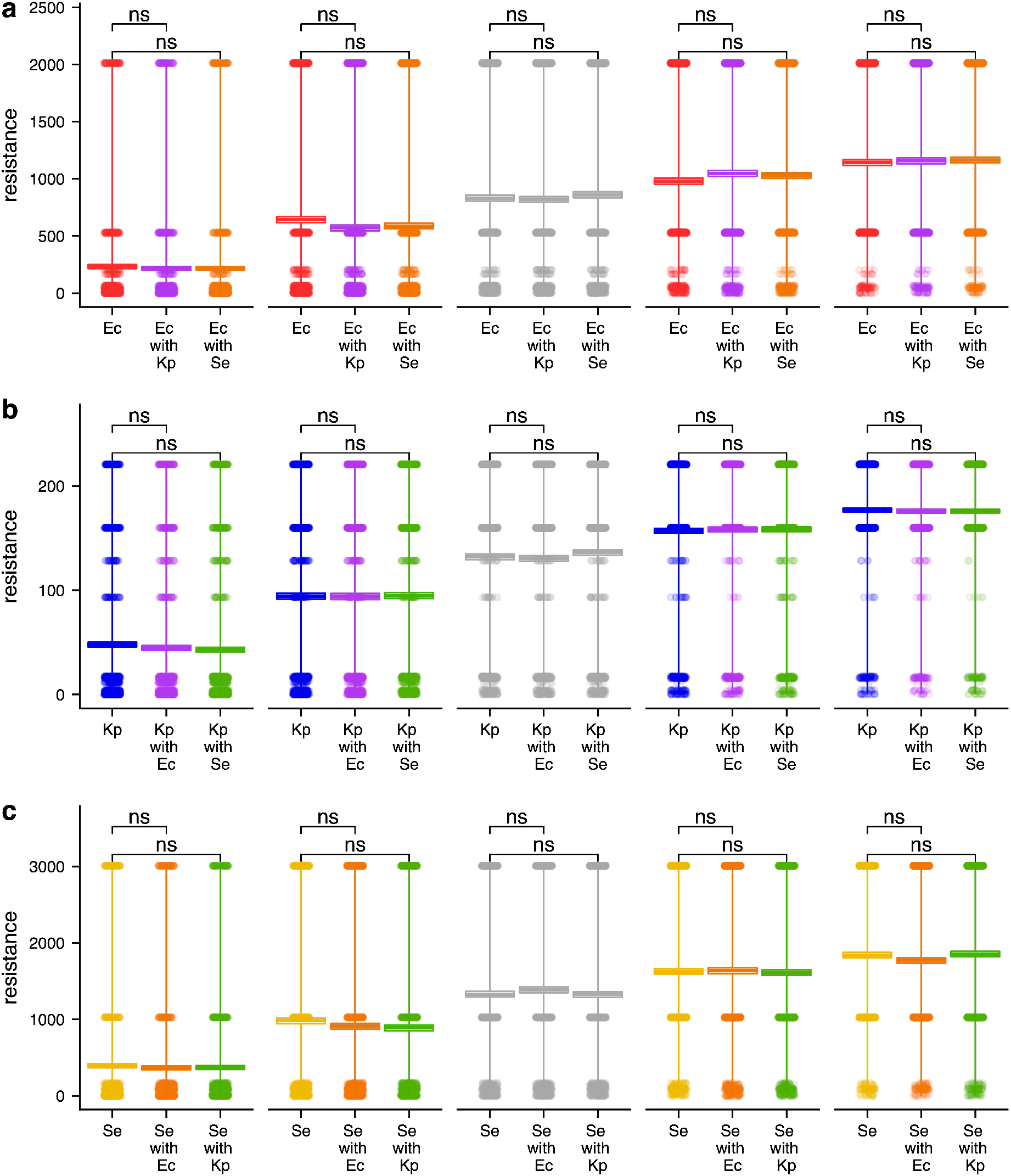
The effect of cumulative time on the evolutionary trends of the *bla* gene. The graphical representation is the same as SI Figure 2. The cumulative time increases from left to right (30, 45, 60, 75, and 90). The gray triplicate in each part indicates the baseline simulation given in SI Figure 2.

**SI Figure 8:**
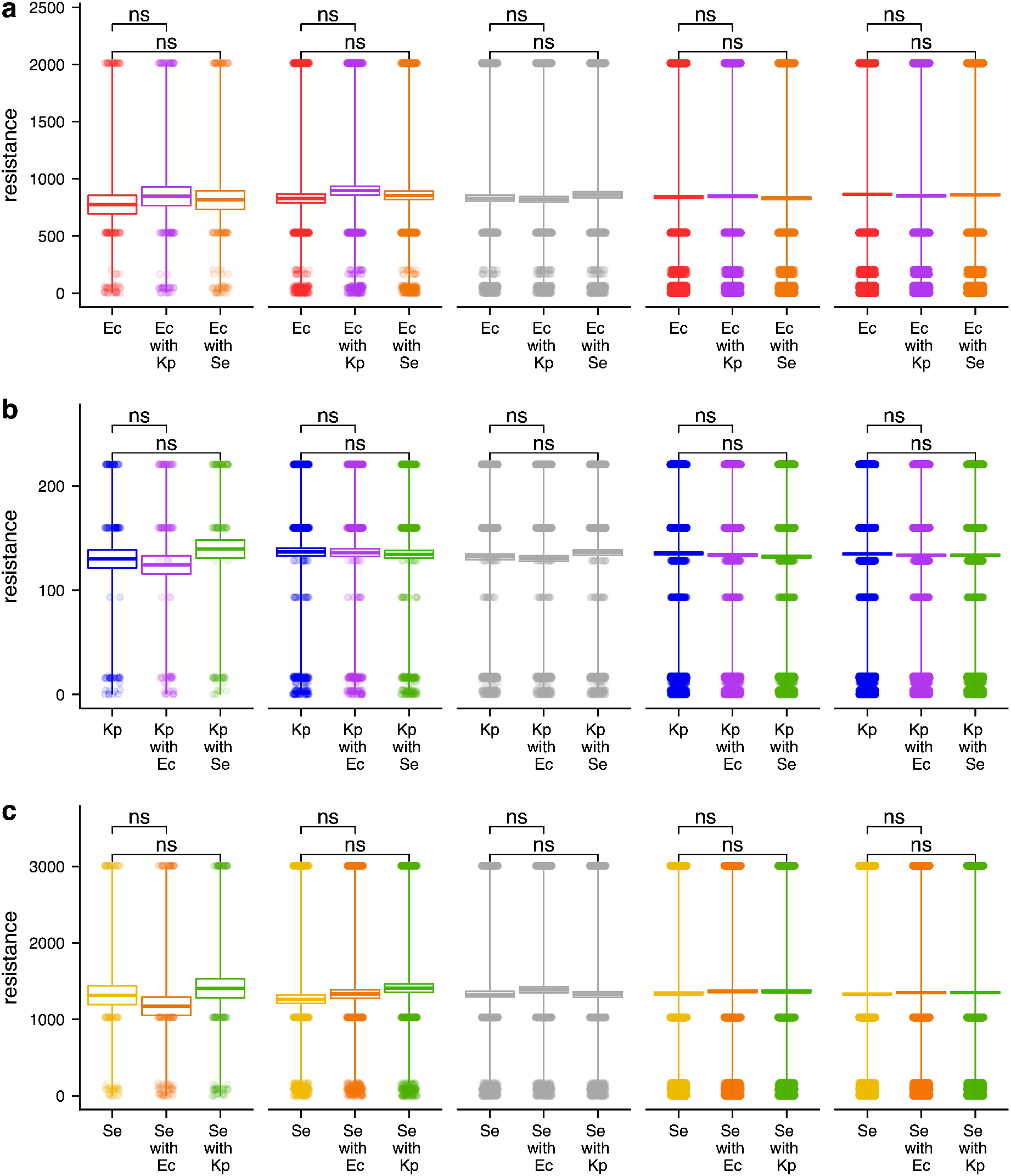
The effect of simulation replicates on the evolutionary trends of the *bla* gene. The graphical representation is the same as SI Figure 2. The cumulative time increases from left to right (100, 500, 1000, 5000, and 10000). The gray triplicate in each part indicates the baseline simulation given in SI Figure 2.

**SI Figure 9:**
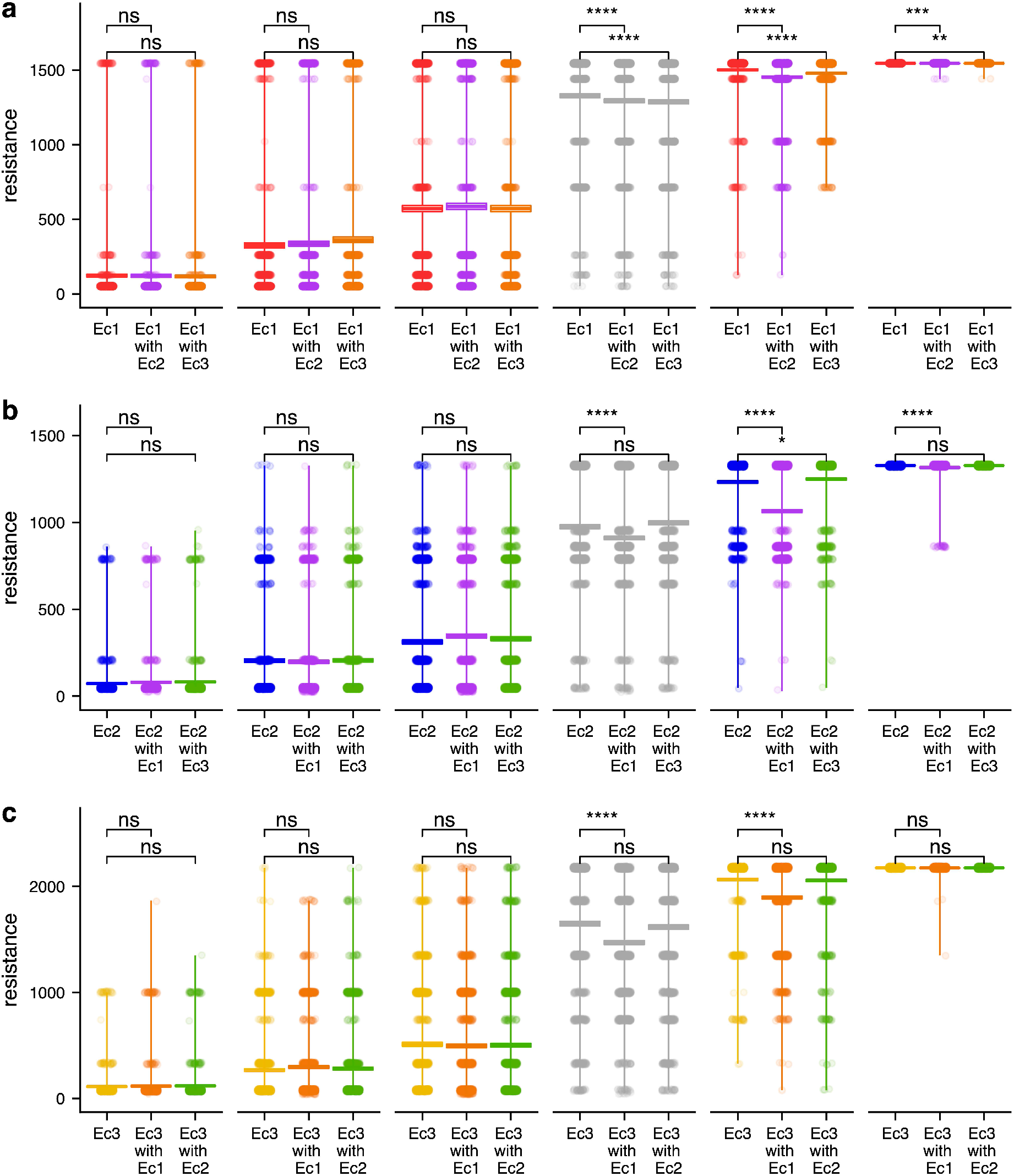
The effect of mutation rates on the evolutionary trends of the DHFR gene. The graphical representation is the same as SI Figure 2. The mutation rate increases from left to right (1 × 10^6^, 5 × 10^6^, 1 × 10^5^, 5 × 10^5^, 1 × 10^4^, and 5 × 10^4^). The gray triplicate in each part indicates the baseline simulation given in SI Figure 5.

**SI Figure 10:**
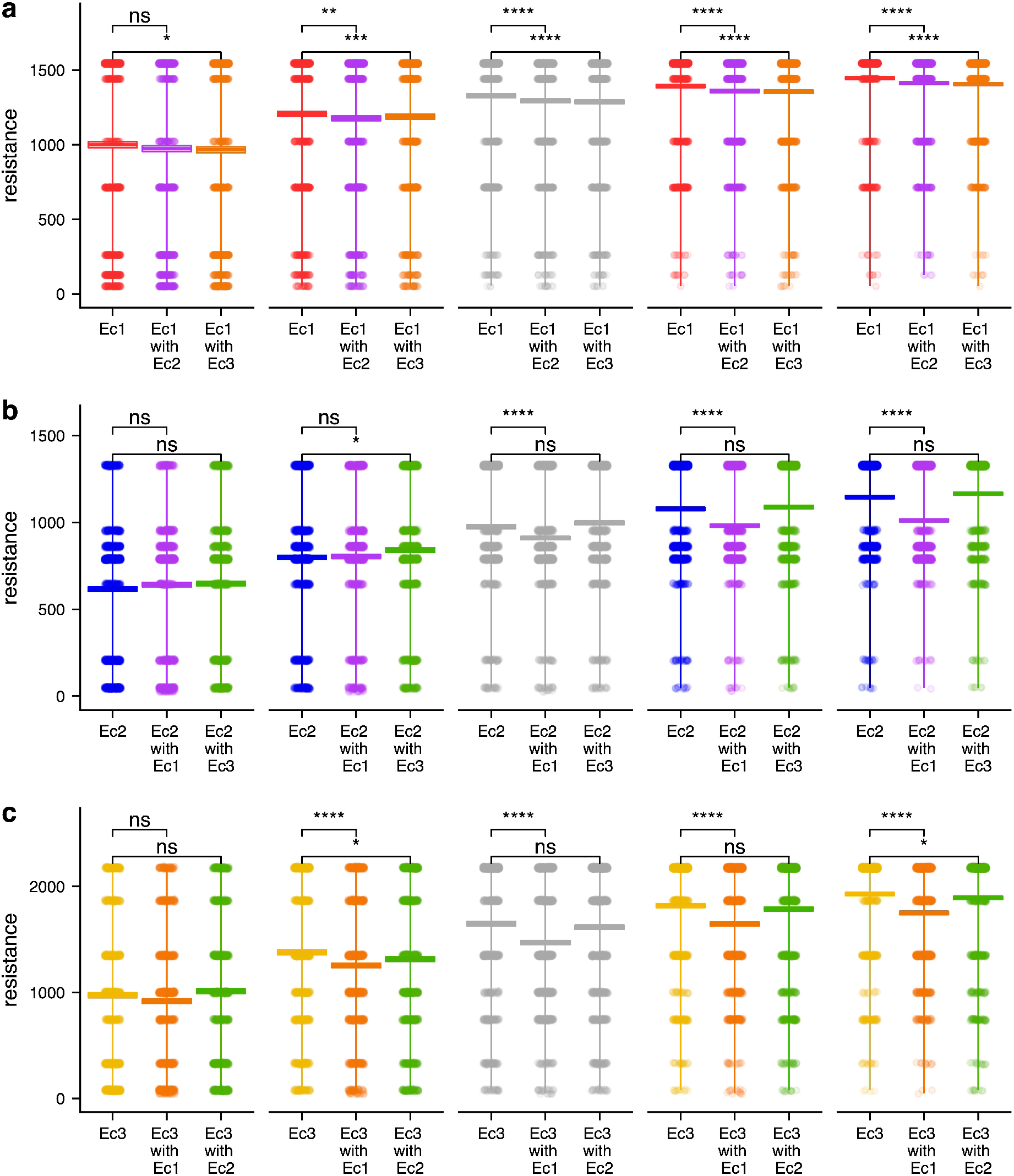
The effect of cumulative time on the evolutionary trends of the DHFR gene. The graphical representation is the same as SI Figure 2. The cumulative time increases from left to right (18, 27, 36, 45, and 54). The gray triplicate in each part indicates the baseline simulation given in SI Figure 5.

**SI Figure 11:**
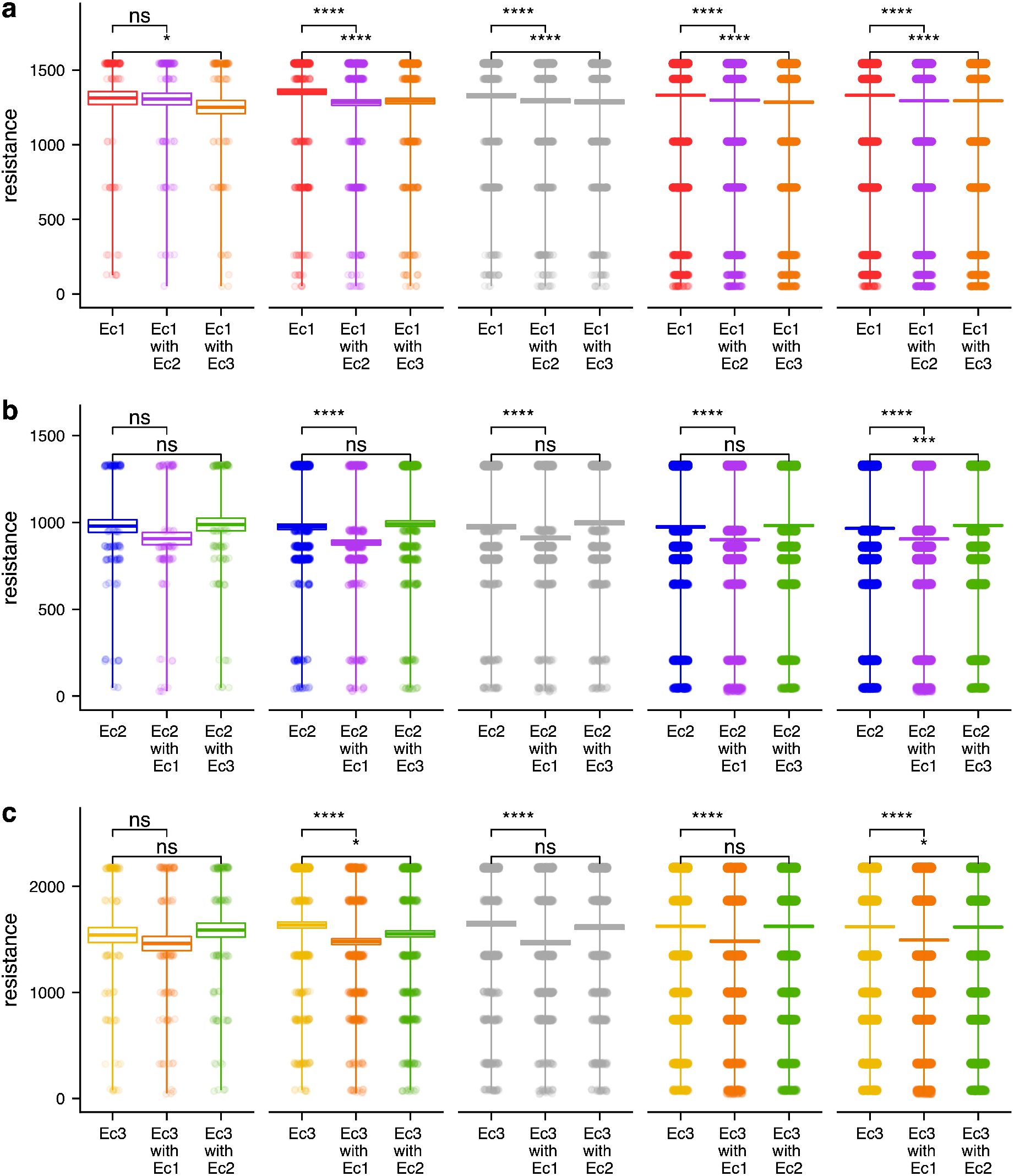
The effect of simulation replicates on the evolutionary trends of the DHFR gene. The graphical representation is the same as SI Figure 2. The cumulative time increases from left to right (100, 500, 1000, 5000, and 10000). The gray triplicate in each part indicates the baseline simulation given in SI Figure 5.

**SI Table 1:**
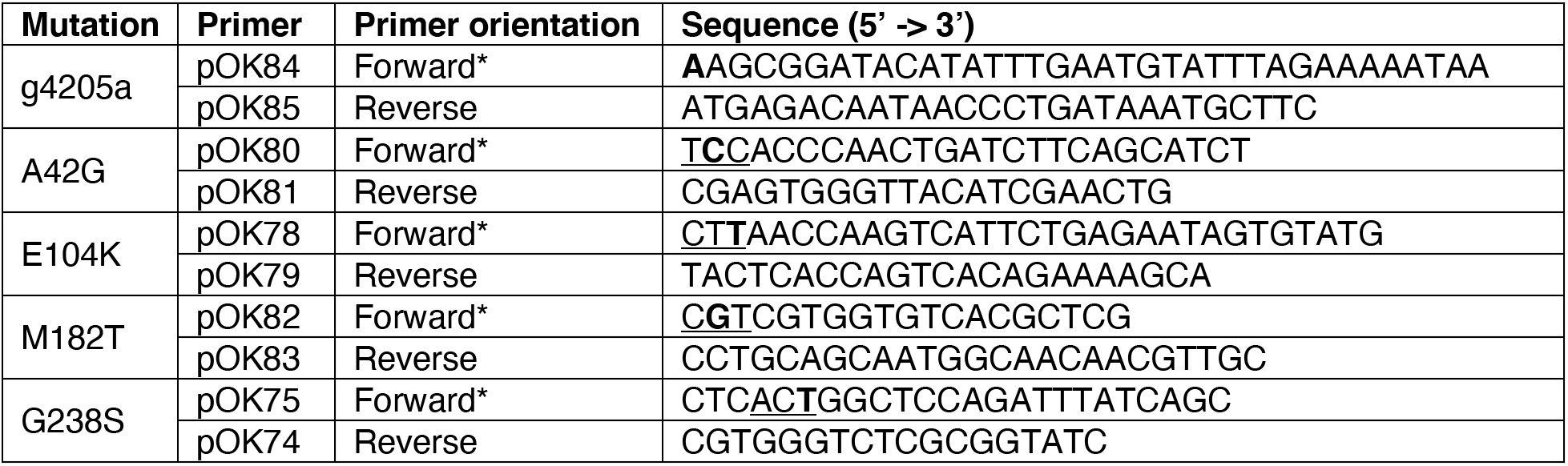
Primers used for Site-Directed Mutagenesis. The mutagenic primer is labelled with an asterisk. If an amino acid is being mutated, the codon is <u>underlined</u>. The nucleotide being mutated is **bolded**.

**SI Table 2 :**
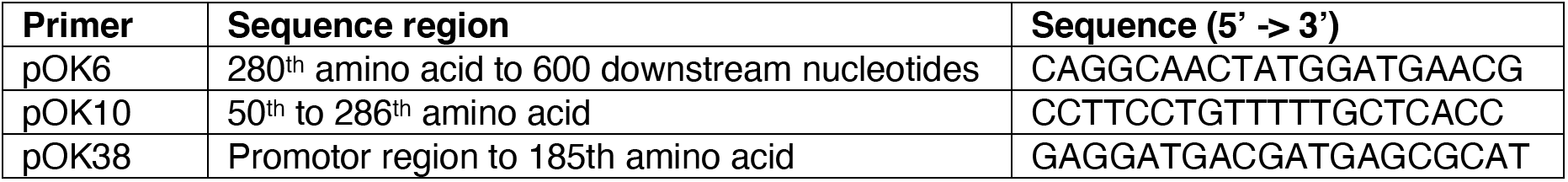
Primers used for Sanger sequencing.

**SI Table 3 :** Engineered variants using Site-Directed Mutagenesis.

| Variant | Number of mutations |
| --- | --- |
| g4205a | 1 |
| A42G | 1 |
| E104K | 1 |
| M182T | 1 |
| G238S | 1 |
| g4205a, A42G | 2 |
| g4205a, E104K | 2 |
| g4205a, M182T | 2 |
| g4205a, G238S | 2 |
| A42G, E104K | 2 |
| A42G, M182T | 2 |
| A42G, G238S | 2 |
| E104K, M182T | 2 |
| E104K, G238S | 2 |
| M182T, G238S | 2 |
| g4205a, A42G, E104K | 3 |
| g4205a, A42G, M182T | 3 |
| g4205a, A42G, G238S | 3 |
| g4205a, E104K, M182T | 3 |
| g4205a, E104K, G238S | 3 |
| g4205a, M182T, G238S | 3 |
| A42G, E104K, M182T | 3 |
| A42G, E104K, G238S | 3 |
| A42G, M182T, G238S | 3 |
| E104K, M182T, G238S | 3 |
| g4205a, A42G, E104K, M182T | 4 |
| g4205a, A42G, E104K, G238S | 4 |
| g4205a, A42G, M182T, G238S | 4 |
| g4205a, E104K, M182T, G238S | 4 |
| A42G, E104K, M182T, G238S | 4 |
| g4205a, A42G, E104K, M182T, G238S | 5 |

**SI Table 4 :**
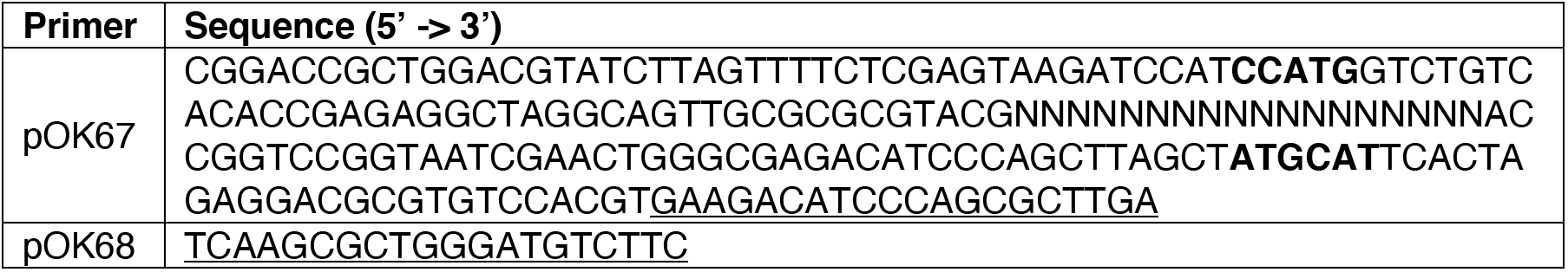
Primers used for creating the barcode fragment. The NcoI and NsiI restriction sites are **bolded.** The homologous nucleotides used for creating the double > stranded fragment are <u>underlined</u>.

**SI Table 5 :** Initial cell density of each library before selection in CTX.

| Species | Number of replicates | Average initial density (cfu ml<sup>-1</sup>) | Standard error |
| --- | --- | --- | --- |
| **Ec** | 6 | 4.07 x 10<sup>5</sup> | 2.67 x 10<sup>4</sup> |
| **Kp** | 6 | 3.21 x 10<sup>5</sup> | 2.99 x 10<sup>4</sup> |
| **Se** | 6 | 6.07 x 10<sup>5</sup> | 3.41 x 10<sup>4</sup> |

**SI Table 6 :** Final cell density (cfu ml^-1^) from each library selection. The concentrations where sequencing data was obtained are **bolded** for each species. A portion of the lower concentration was not submitted for sequencing given the resistance level of the ancestral allele, TEM-1. Test tubes that were not turbid (NT) after the 24 h incubation are designated.

| CTX concentration (µg ml<sup>-1</sup>) | **Ec** | **Kp** | **Se** |
| --- | --- | --- | --- |
| 0.0000 | **2.44 x 10<sup>9</sup>** | **3.00 x 10<sup>9</sup>** | **2.84 x 10<sup>9</sup>** |
| 0.00393 | 2.28 x 10<sup>9</sup> | 3.88 x 10<sup>9</sup> | 1.91 x 10<sup>9</sup> |
| 0.0056 | 2.32 x 10<sup>9</sup> | 3.32 x 10<sup>9</sup> | 2.26 x 10<sup>9</sup> |
| 0.0079 | 1.72 x 10<sup>9</sup> | 3.52 x 10<sup>9</sup> | 2.39 x 10<sup>9</sup> |
| 0.0112 | 2.00 x 10<sup>9</sup> | 3.22 x 10<sup>9</sup> | 2.33 x 10<sup>9</sup> |
| 0.0158 | 1.42 x 10<sup>9</sup> | 2.24 x 10<sup>9</sup> | 2.15 x 10<sup>9</sup> |
| 0.0223 | 1.06 x 10<sup>9</sup> | 2.88 x 10<sup>9</sup> | 1.97 x 10<sup>9</sup> |
| 0.0315 | **1.06 x 10<sup>9</sup>** | **2.36 x 10<sup>9</sup>** | **2.37 x 10<sup>9</sup>** |
| 0.0445 | **9.80 x 10<sup>8</sup>** | **2.18 x 10<sup>9</sup>** | **1.70 x 10<sup>9</sup>** |
| 0.0629 | **9.80 x 10<sup>8</sup>** | **3.48 x 10<sup>9</sup>** | **1.92 x 10<sup>9</sup>** |
| 0.0889 | **1.06 x 10<sup>9</sup>** | **3.04 x 10<sup>9</sup>** | **2.01 x 10<sup>9</sup>** |
| 0.1257 | **1.32 x 10<sup>9</sup>** | **3.44 x 10<sup>9</sup>** | **1.74 x 10<sup>9</sup>** |
| 0.1777 | **1.32 x 10<sup>9</sup>** | **3.88 x 10<sup>9</sup>** | **1.55 x 10<sup>9</sup>** |
| 0.2513 | **1.46 x 10<sup>9</sup>** | **3.20 x 10<sup>9</sup>** | **2.07 x 10<sup>9</sup>** |
| 0.3553 | **1.26 x 10<sup>9</sup>** | **3.08 x 10<sup>9</sup>** | **1.92 x 10<sup>9</sup>** |
| 0.5024 | **1.50 x 10<sup>9</sup>** | **4.00 x 10<sup>9</sup>** | **1.91 x 10<sup>9</sup>** |
| 0.7104 | **1.24 x 10<sup>9</sup>** | **3.12 x 10<sup>9</sup>** | **1.92 x 10<sup>9</sup>** |
| 1.0045 | **1.86 x 10<sup>9</sup>** | **3.10 x 10<sup>9</sup>** | **2.05 x 10<sup>9</sup>** |
| 1.42 | **1.26 x 10<sup>9</sup>** | **2.78 x 10<sup>9</sup>** | **1.71 x 10<sup>9</sup>** |
| 2.01 | **1.32 x 10<sup>9</sup>** | **3.04 x 10<sup>9</sup>** | **2.74 x 10<sup>9</sup>** |
| 2.84 | **7.50 x 10<sup>8</sup>** | **3.76 x 10<sup>9</sup>** | **2.02 x 10<sup>9</sup>** |
| 4.02 | **1.07 x 10<sup>9</sup>** | **3.36 x 10<sup>9</sup>** | **1.98 x 10<sup>9</sup>** |
| 5.68 | **1.08 x 10<sup>9</sup>** | **2.68 x 10<sup>9</sup>** | **1.44 x 10<sup>9</sup>** |
| 8.03 | **1.56 x 10<sup>9</sup>** | **3.24 x 10<sup>9</sup>** | **2.72 x 10<sup>9</sup>** |
| 11.35 | **1.42 x 10<sup>9</sup>** | **3.92 x 10<sup>9</sup>** | **1.82 x 10<sup>9</sup>** |
| 16.05 | **2.54 x 10<sup>9</sup>** | **3.40 x 10<sup>9</sup>** | **2.47 x 10<sup>9</sup>** |
| 22.69 | **1.04 x 10<sup>9</sup>** | **4.44 x 10<sup>9</sup>** | **2.08 x 10<sup>9</sup>** |
| 32.08 | **1.46 x 10<sup>9</sup>** | **3.20 x 10<sup>9</sup>** | **1.94 x 10<sup>9</sup>** |
| 45.36 | **1.36 x 10<sup>9</sup>** | **3.68 x 10<sup>9</sup>** | **1.41 x 10<sup>9</sup>** |
| 64.14 | **1.26 x 10<sup>9</sup>** | **2.64 x 10<sup>9</sup>** | **1.55 x 10<sup>9</sup>** |
| 90.69 | **1.37 x 10<sup>9</sup>** | **1.81 x 10<sup>9</sup>** | **1.16 x 10<sup>9</sup>** |
| 128.24 | **4.80 x 10<sup>9</sup>** | **2.63 x 10<sup>9</sup>** | **1.58 x 10<sup>9</sup>** |
| 181.33 | **1.14 x 10<sup>9</sup>** | NT | **7.00 x 10<sup>8</sup>** |
| 256.4 | NT | NT | **1.15 x 10<sup>9</sup>** |
| 362.55 | NT | NT | **1.92 x 10<sup>8</sup>** |
| 512.65 | NT | NT | NT |
| 724.89 | NT | NT | NT |
| 1024.99 | NT | NT | NT |
| 1449.34 | NT | NT | NT |
| 2049.37 | NT | NT | NT |

**SI Table 7 :**
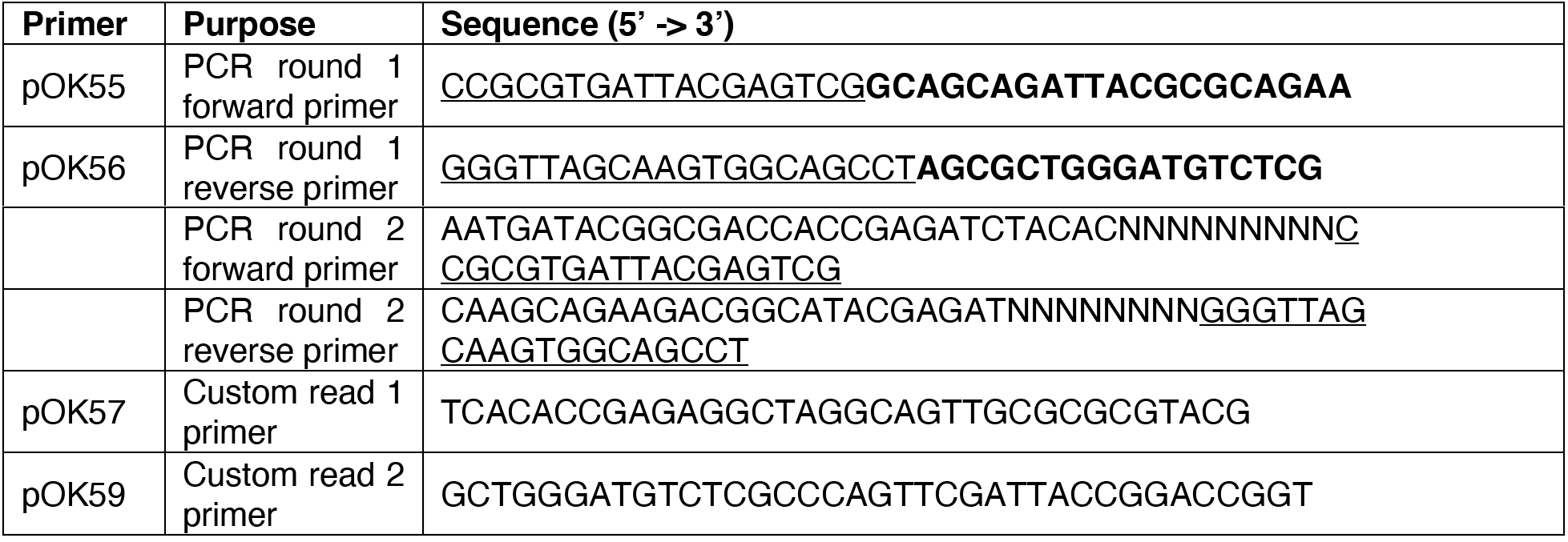
Primers used for library amplification and sequencing. The nucleotides that are homologous to the plasmid are **bolded.** The nucleotides that are homologous to the indexing primers are <u>underlined</u>. The 9bp index used for multiplexing the samples is represented with N nucleotides and are sequence specific depending on the sample.

**SI Table 8 :**
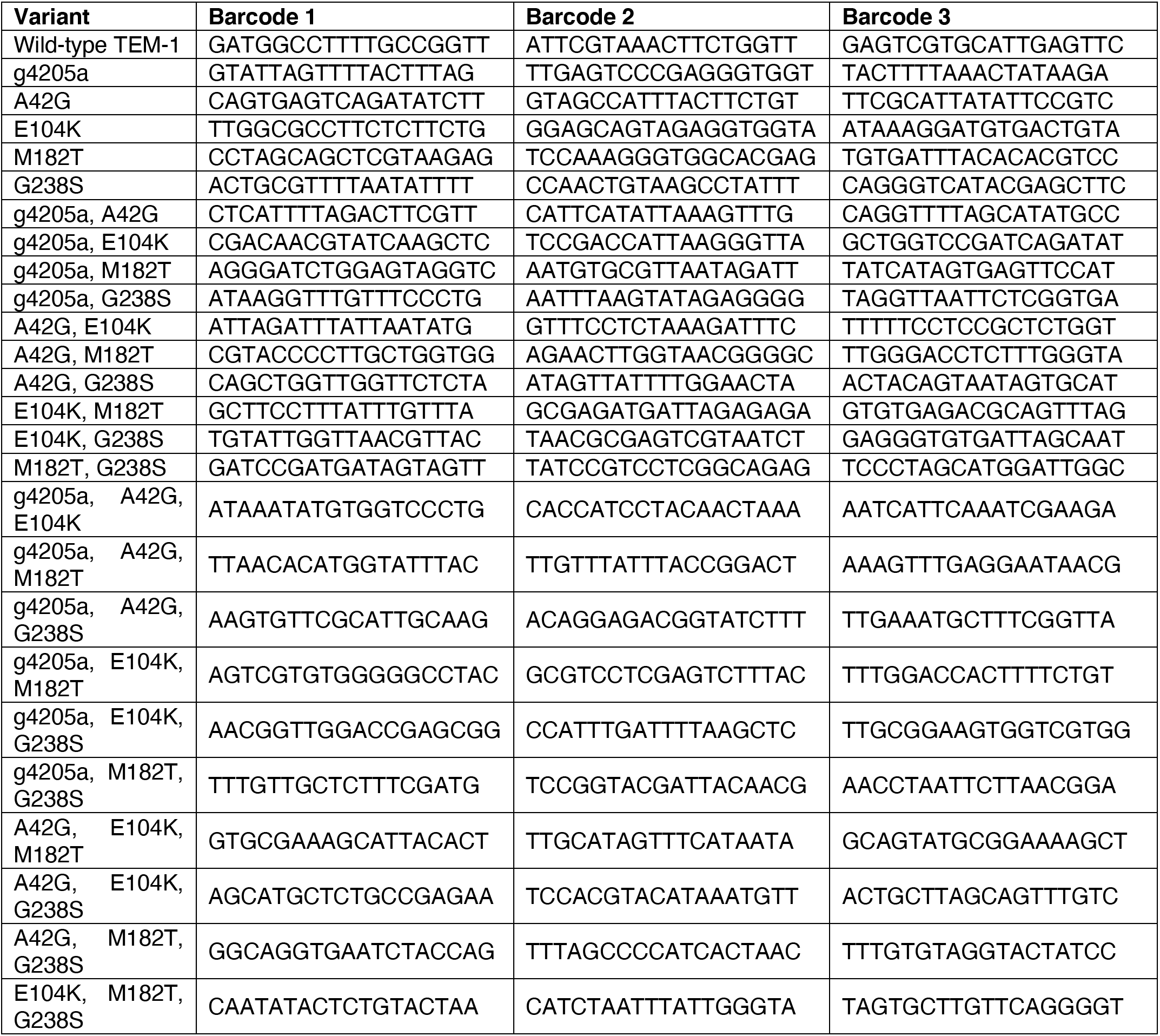
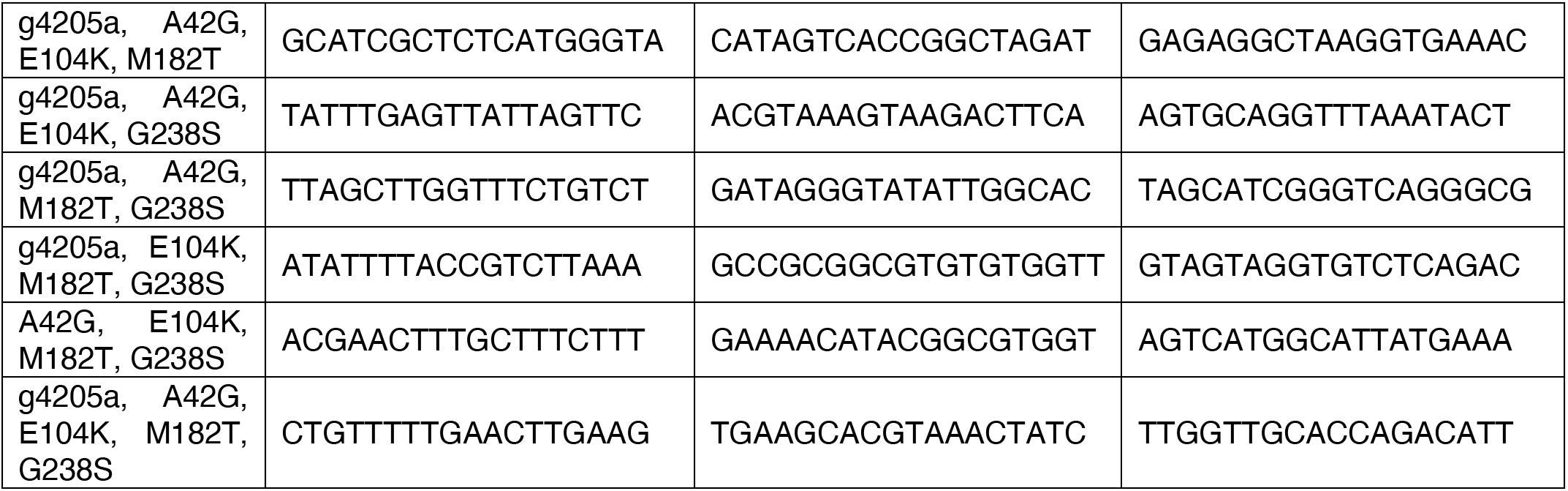
Genotype to barcode map.

**SI Table 9 :**
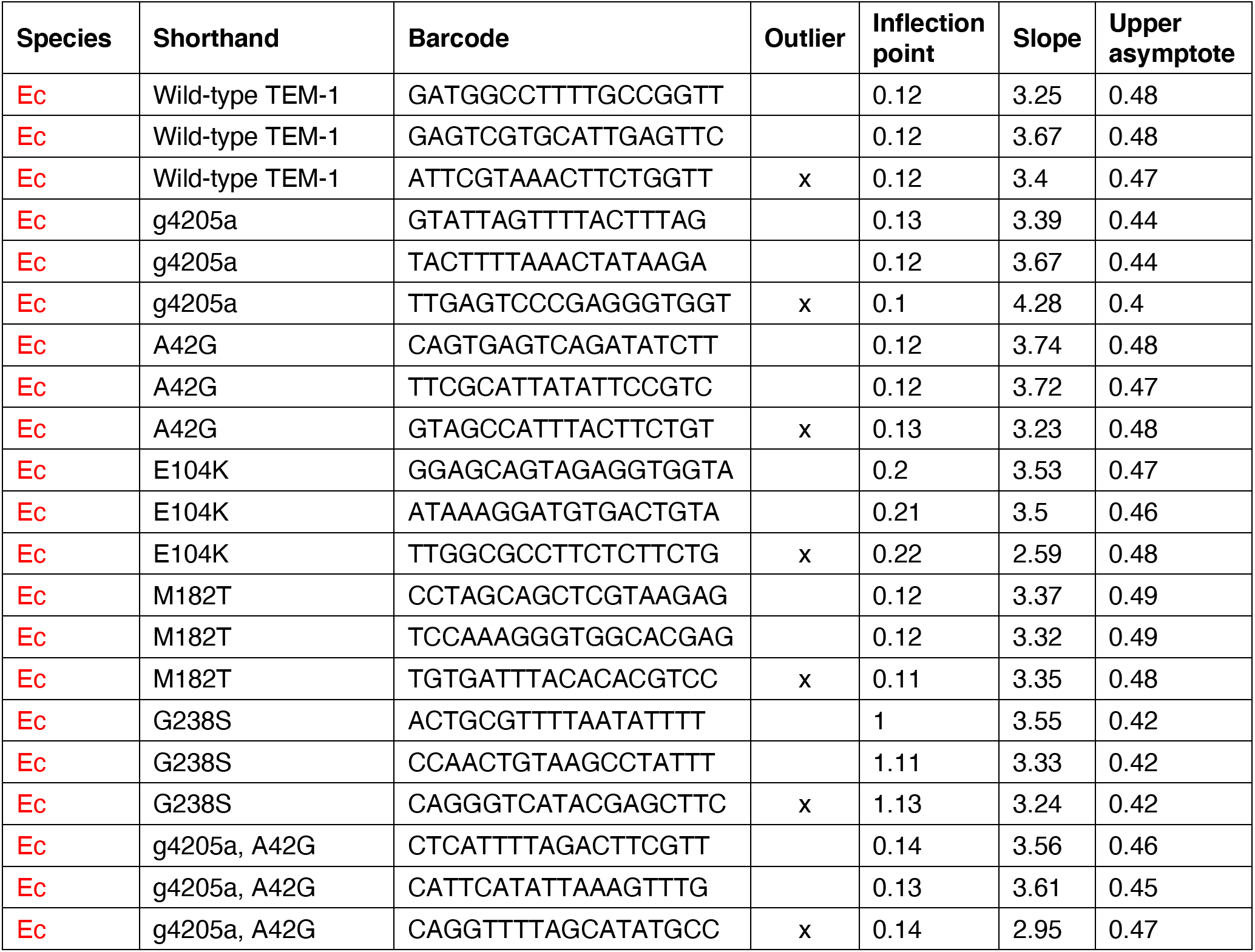
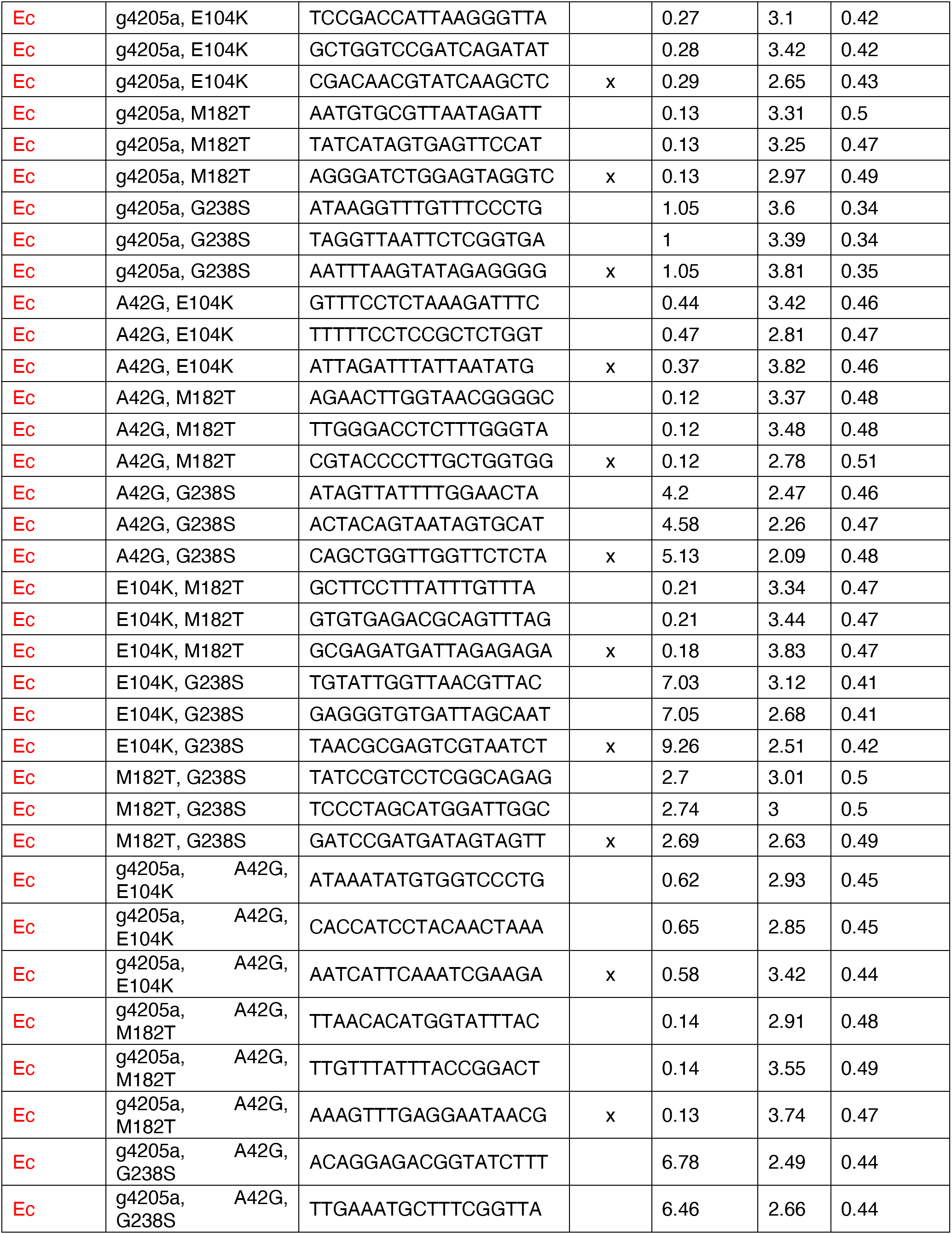
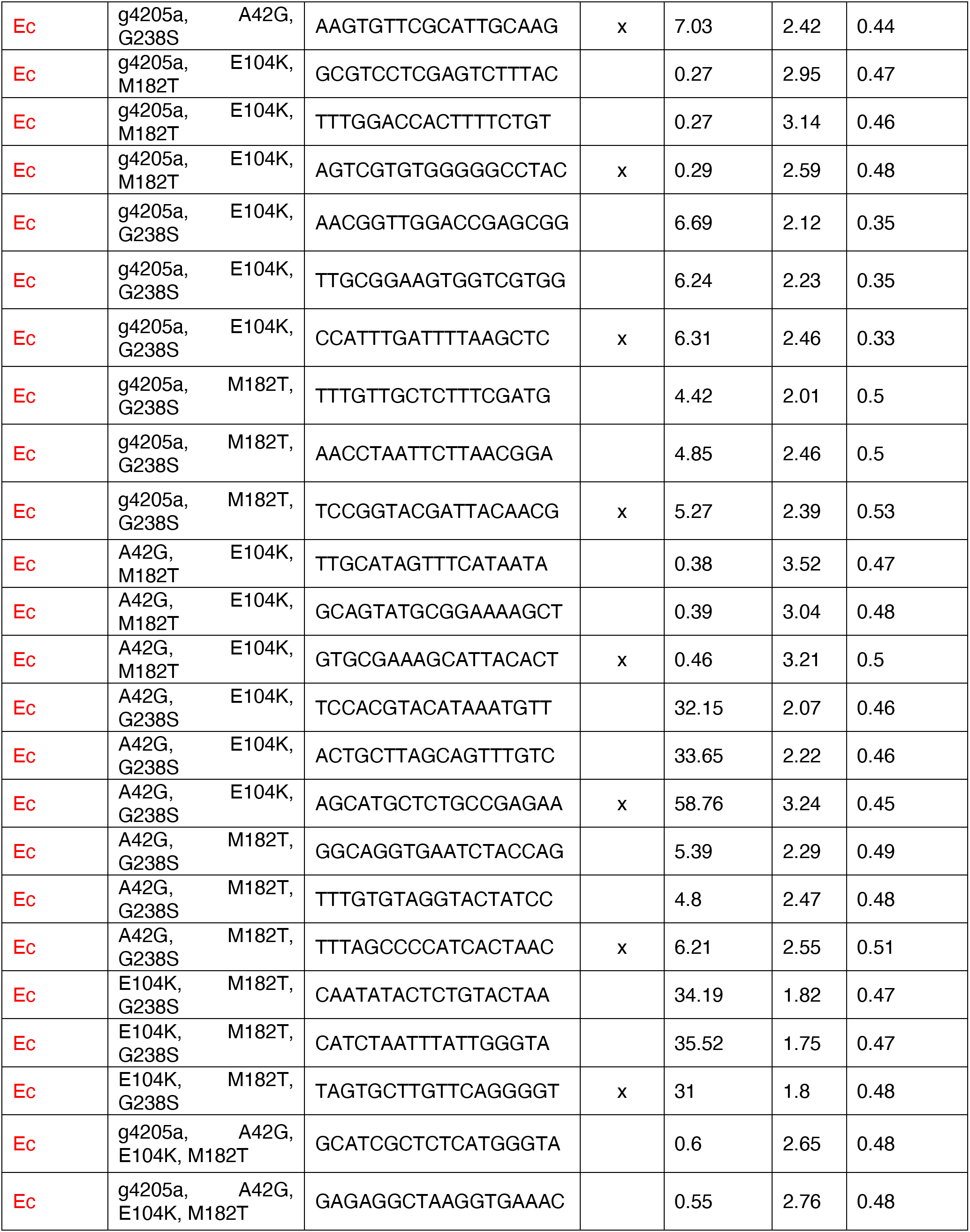
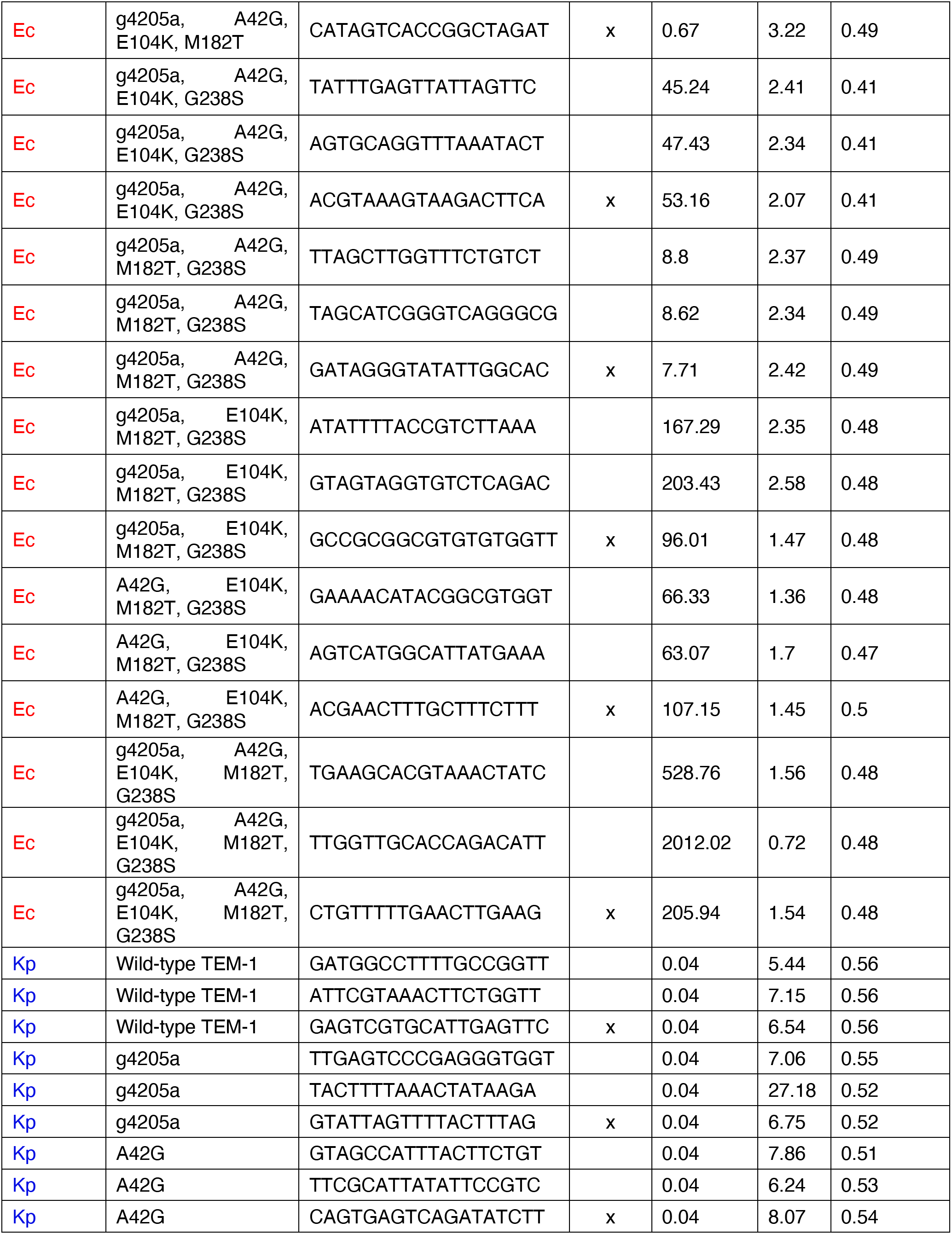
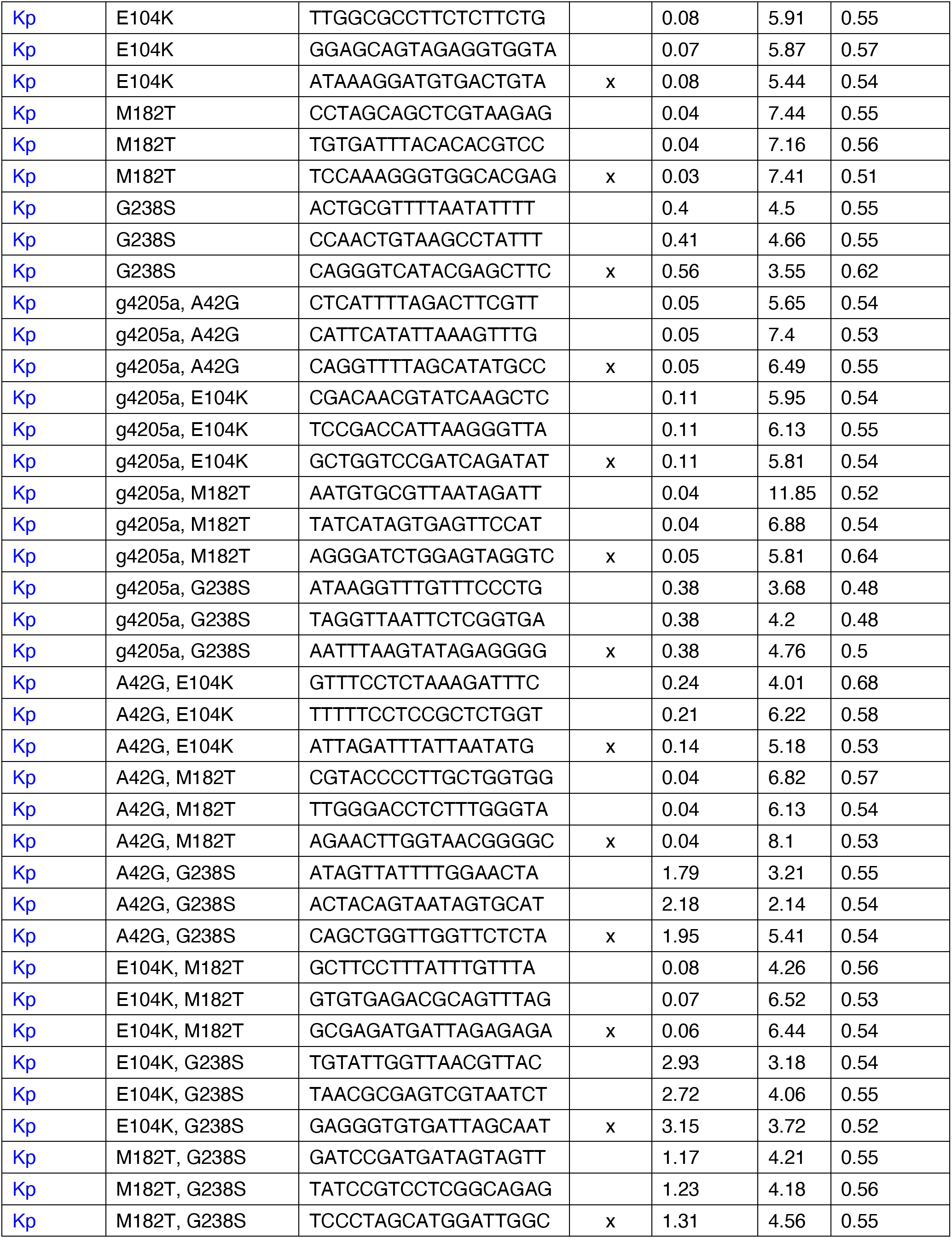
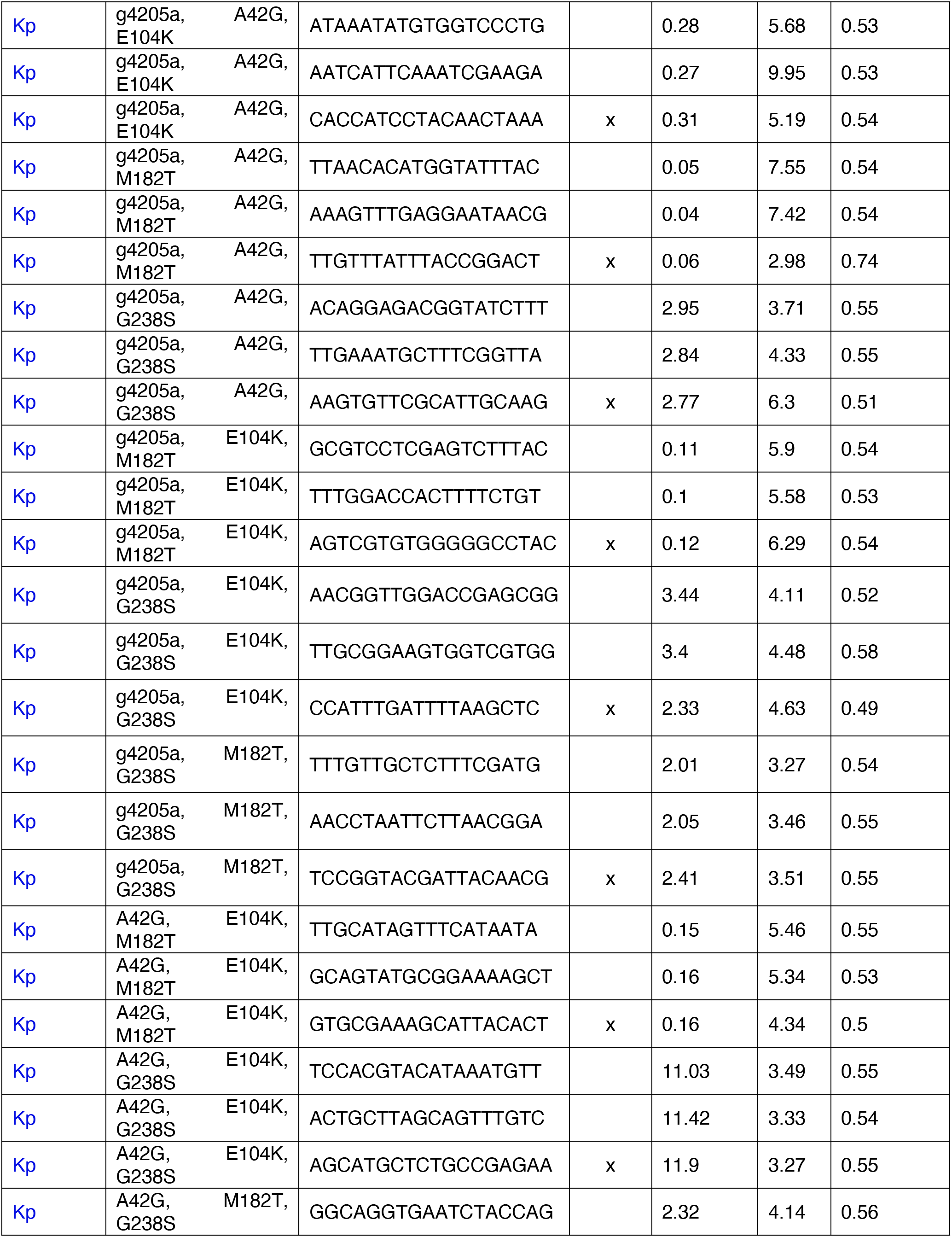
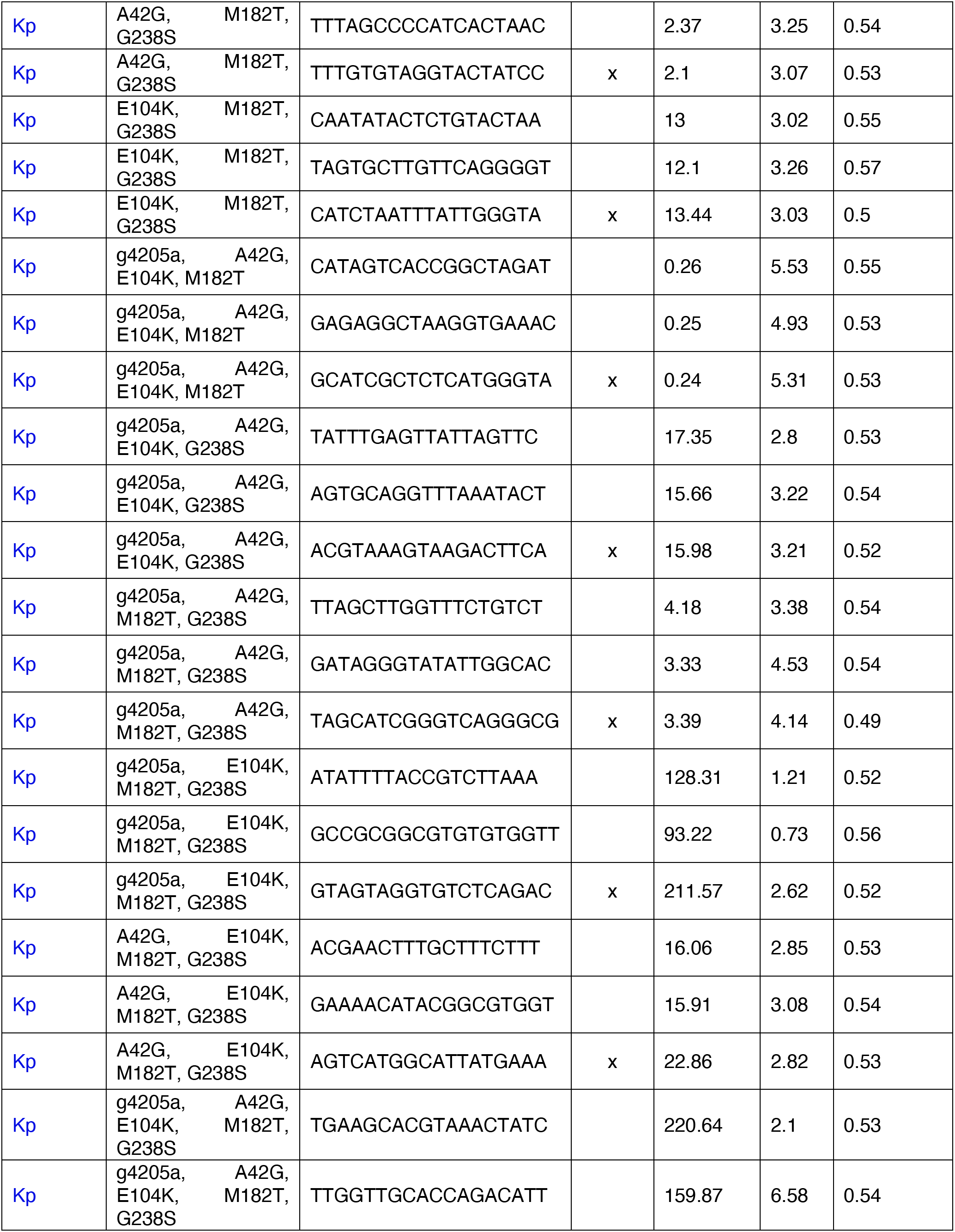
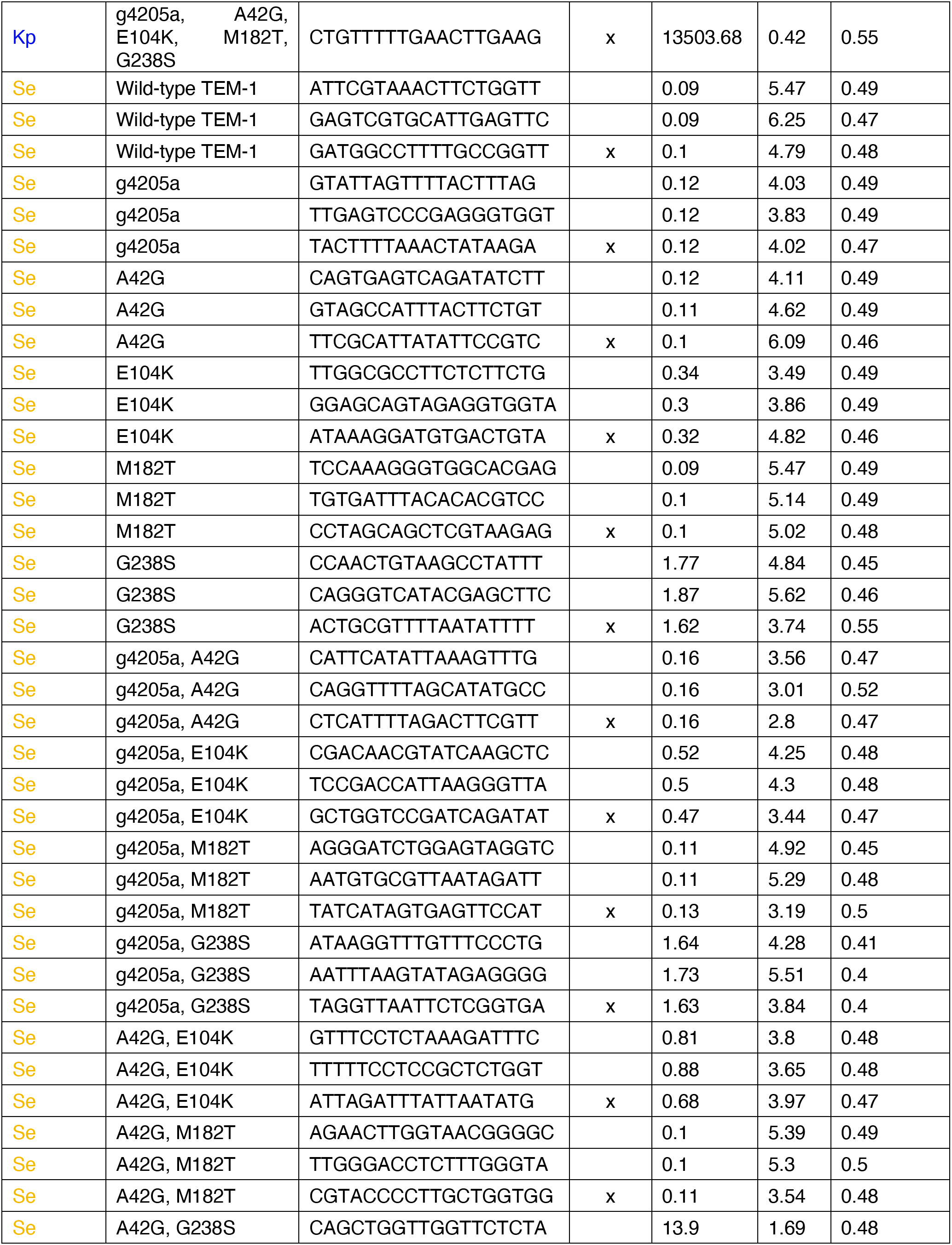
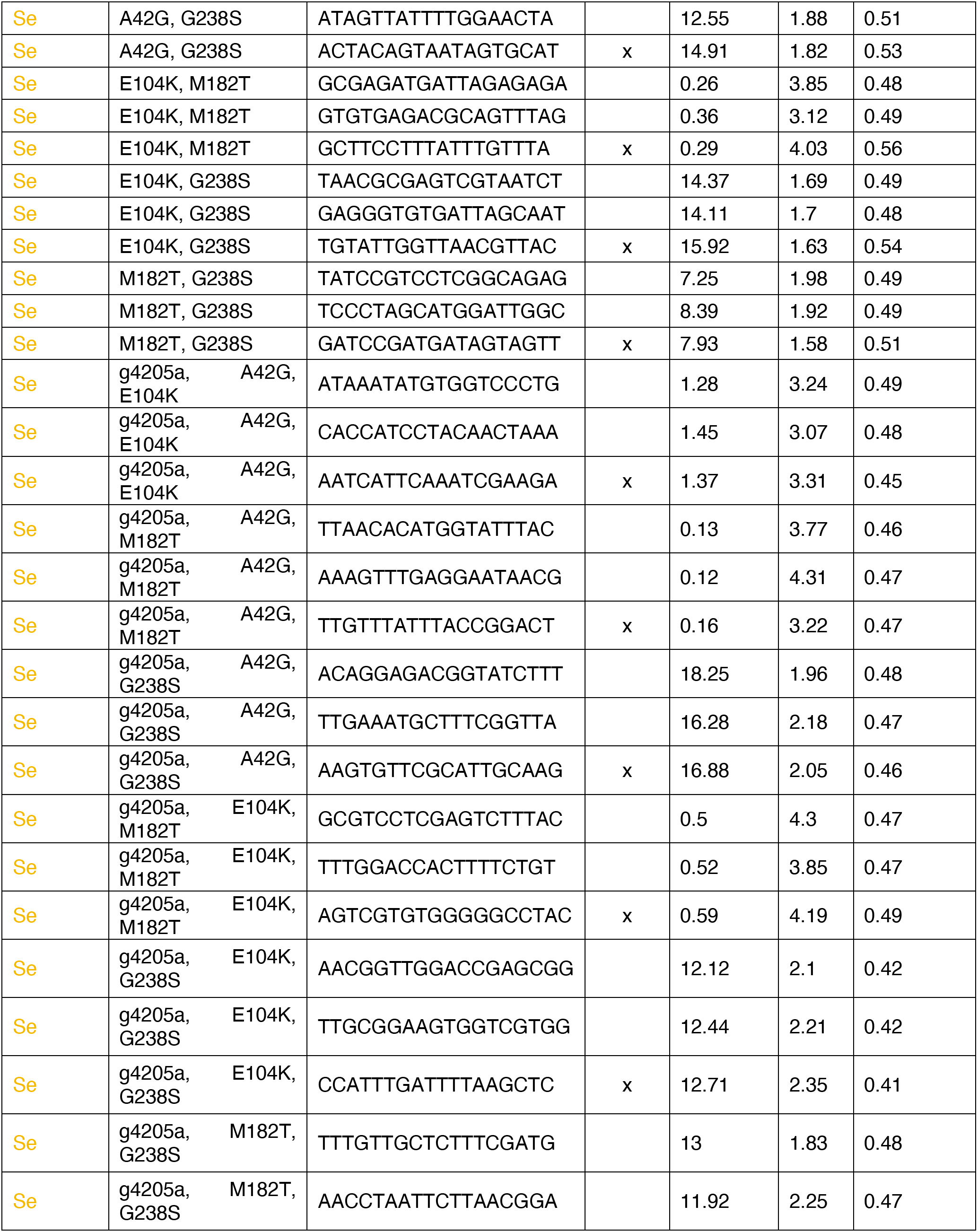
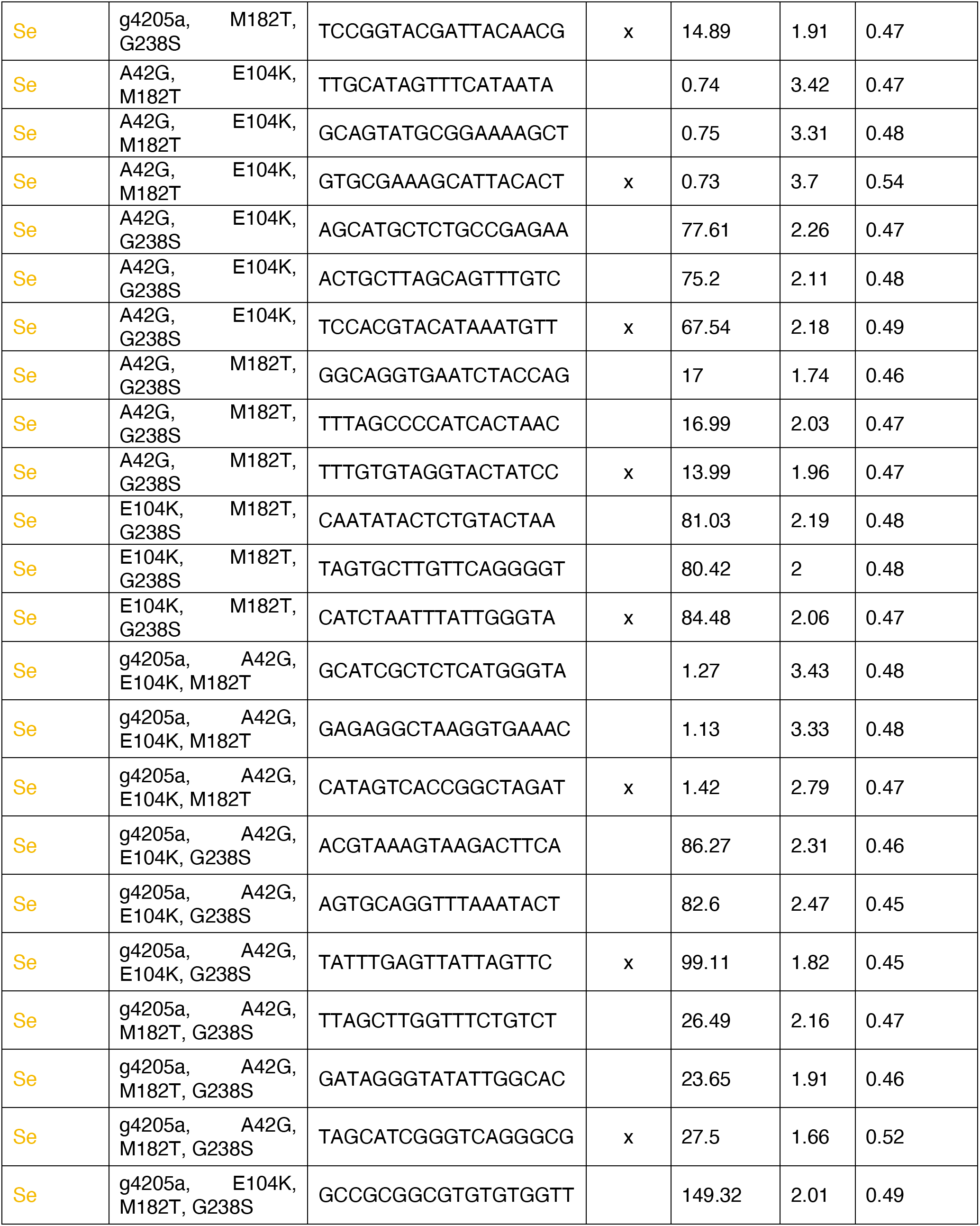
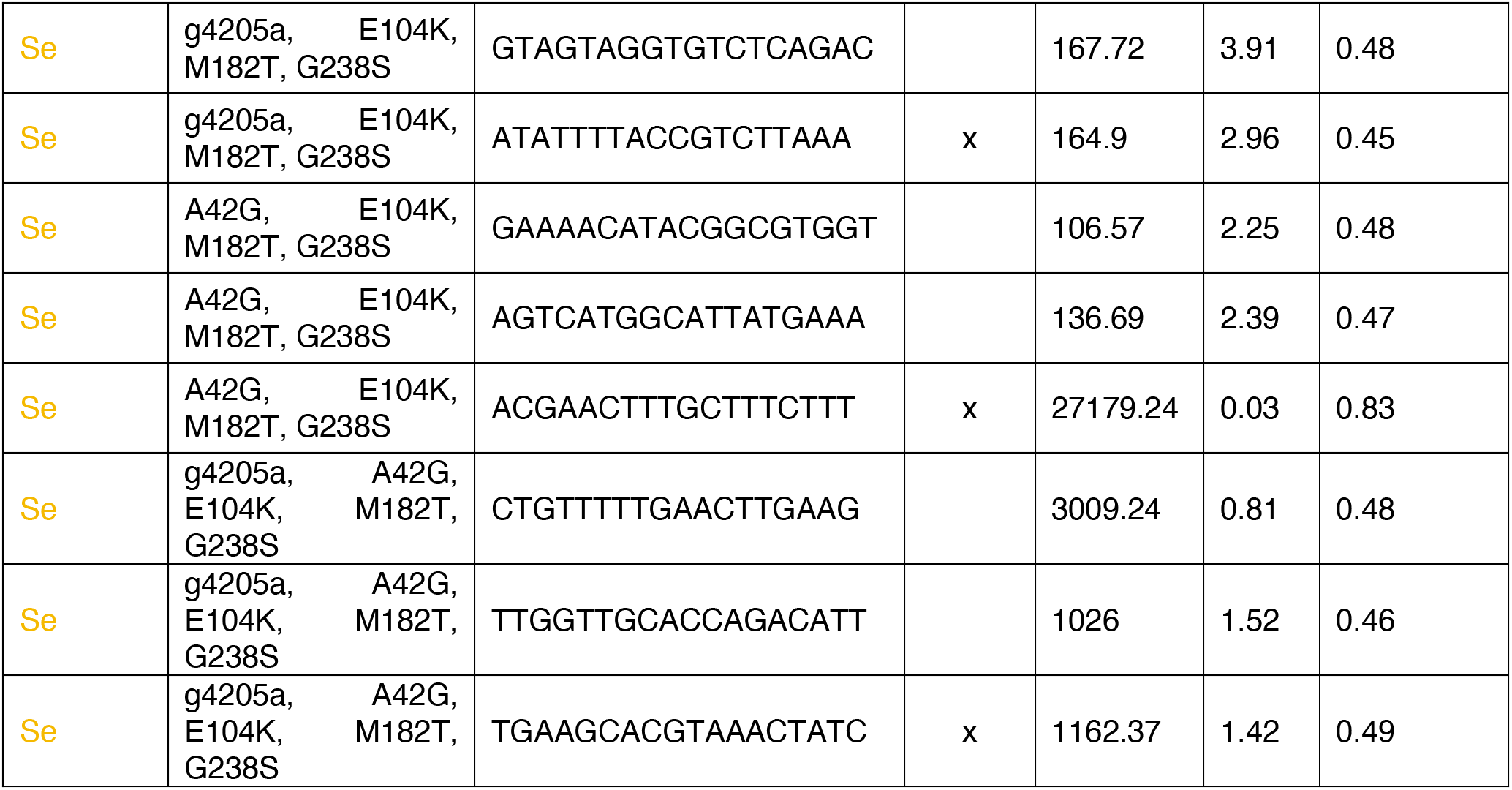
The three-parameter estimates (inflection point, steepness, and upper asymptote) from the dose-response curve fitting for each barcode-allele-species combination.

**SI Table 10 :**
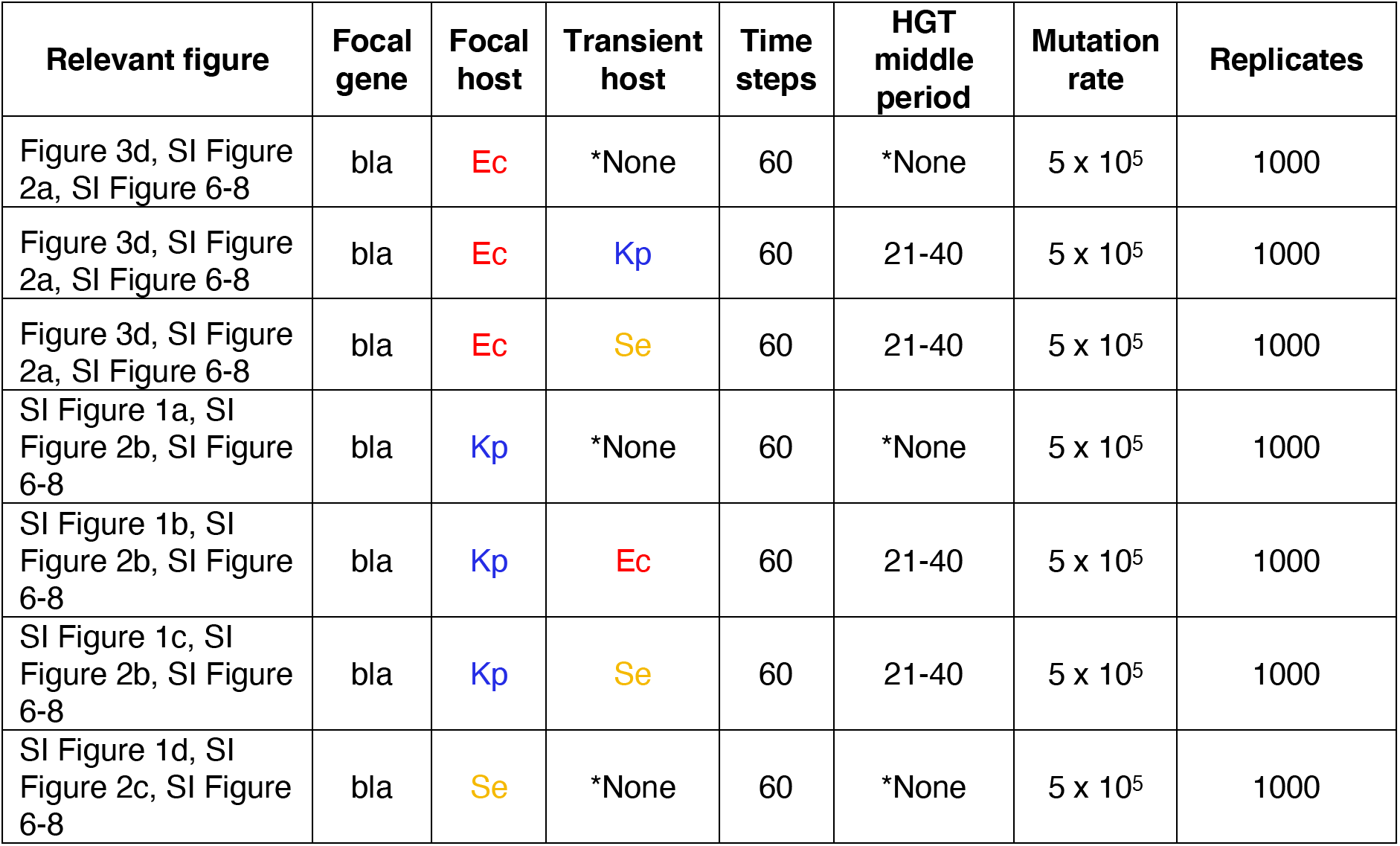
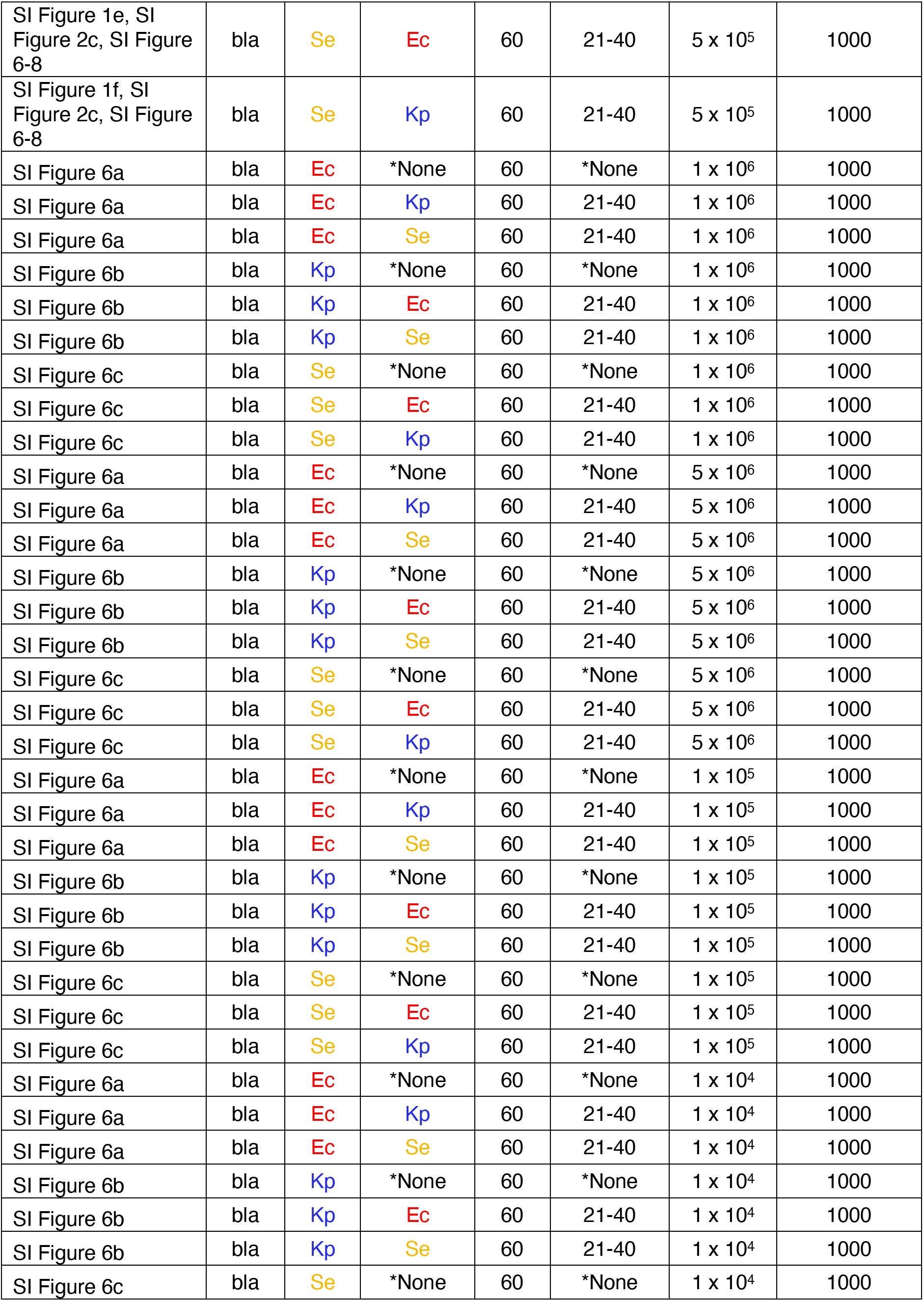
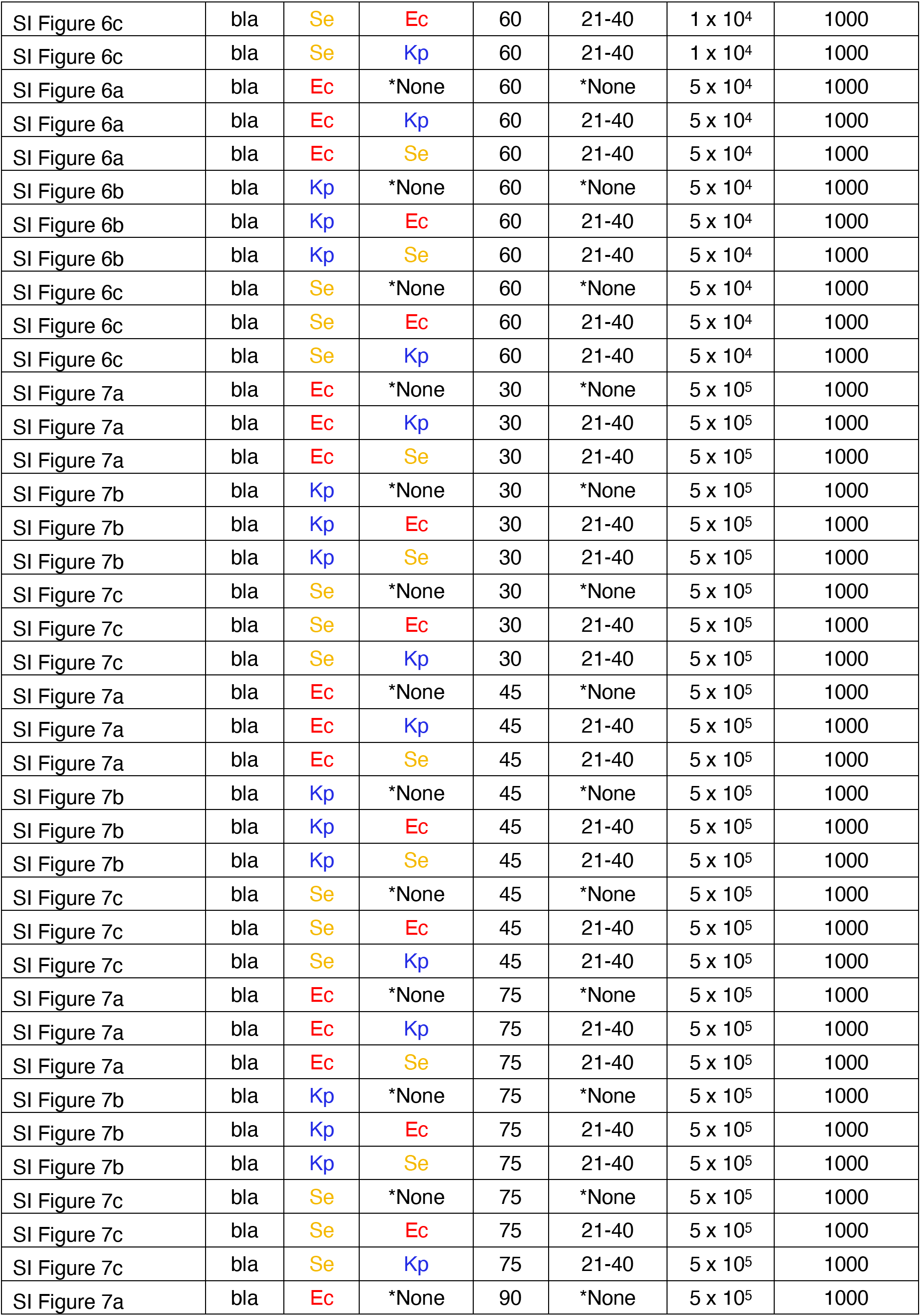
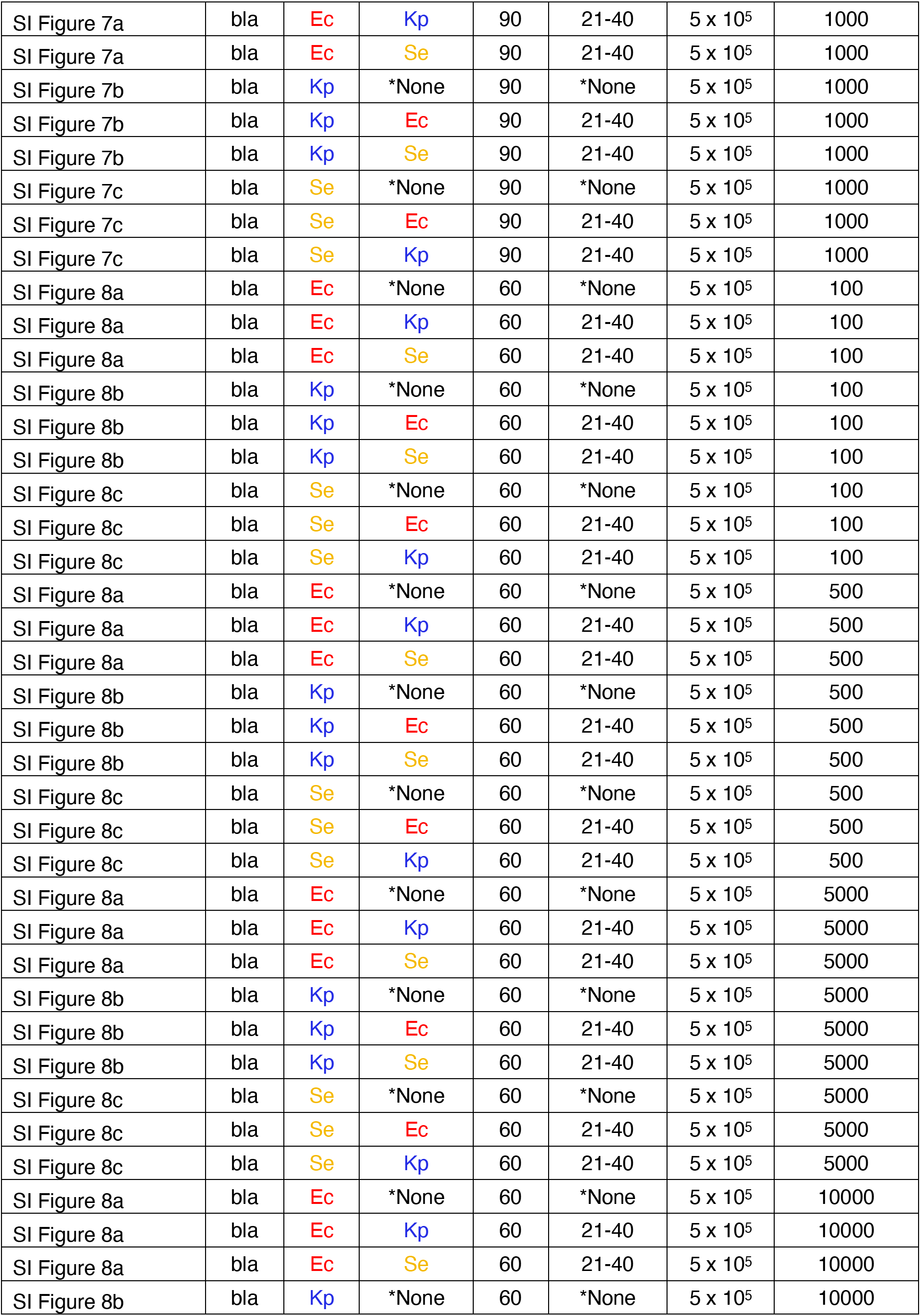
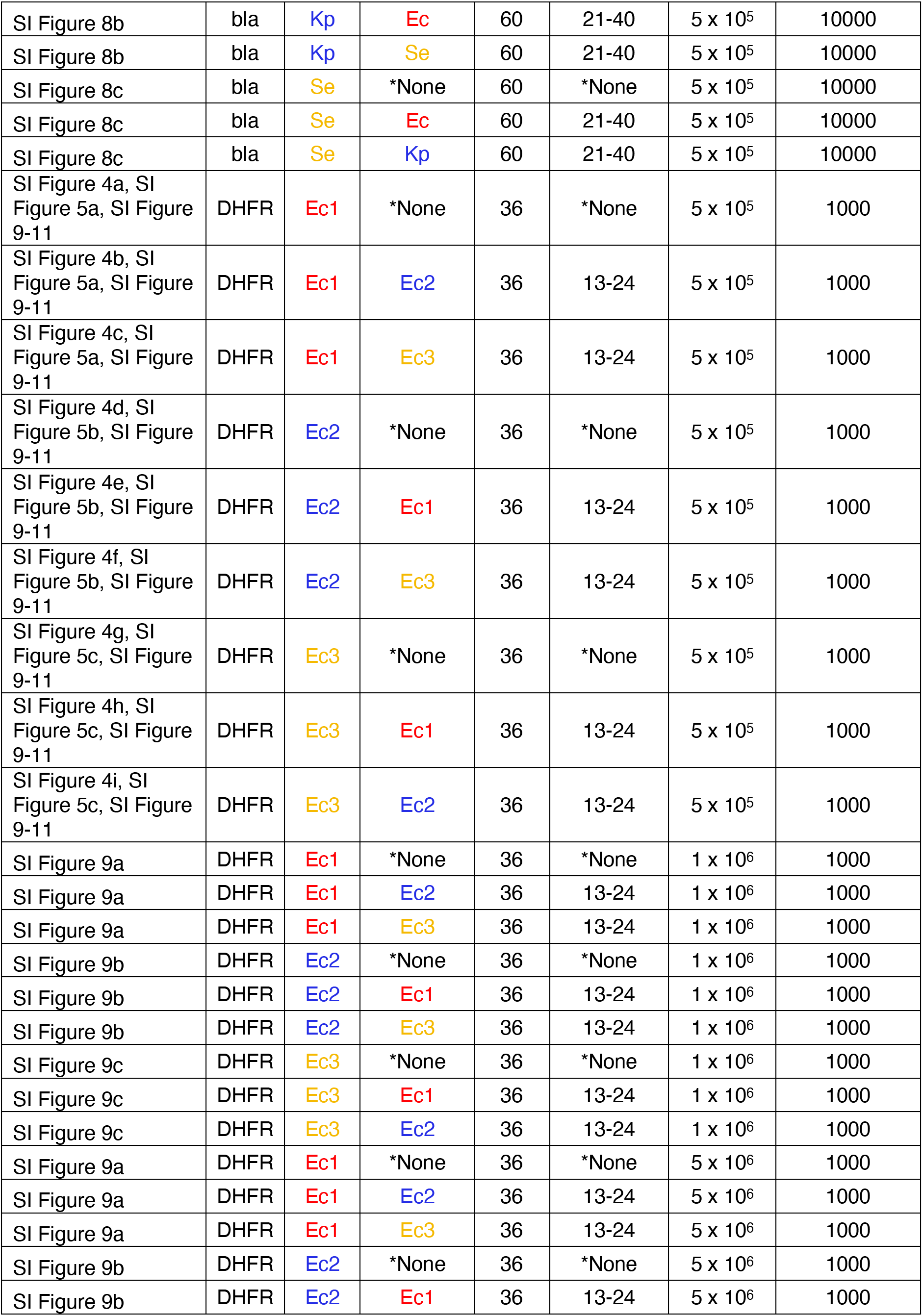
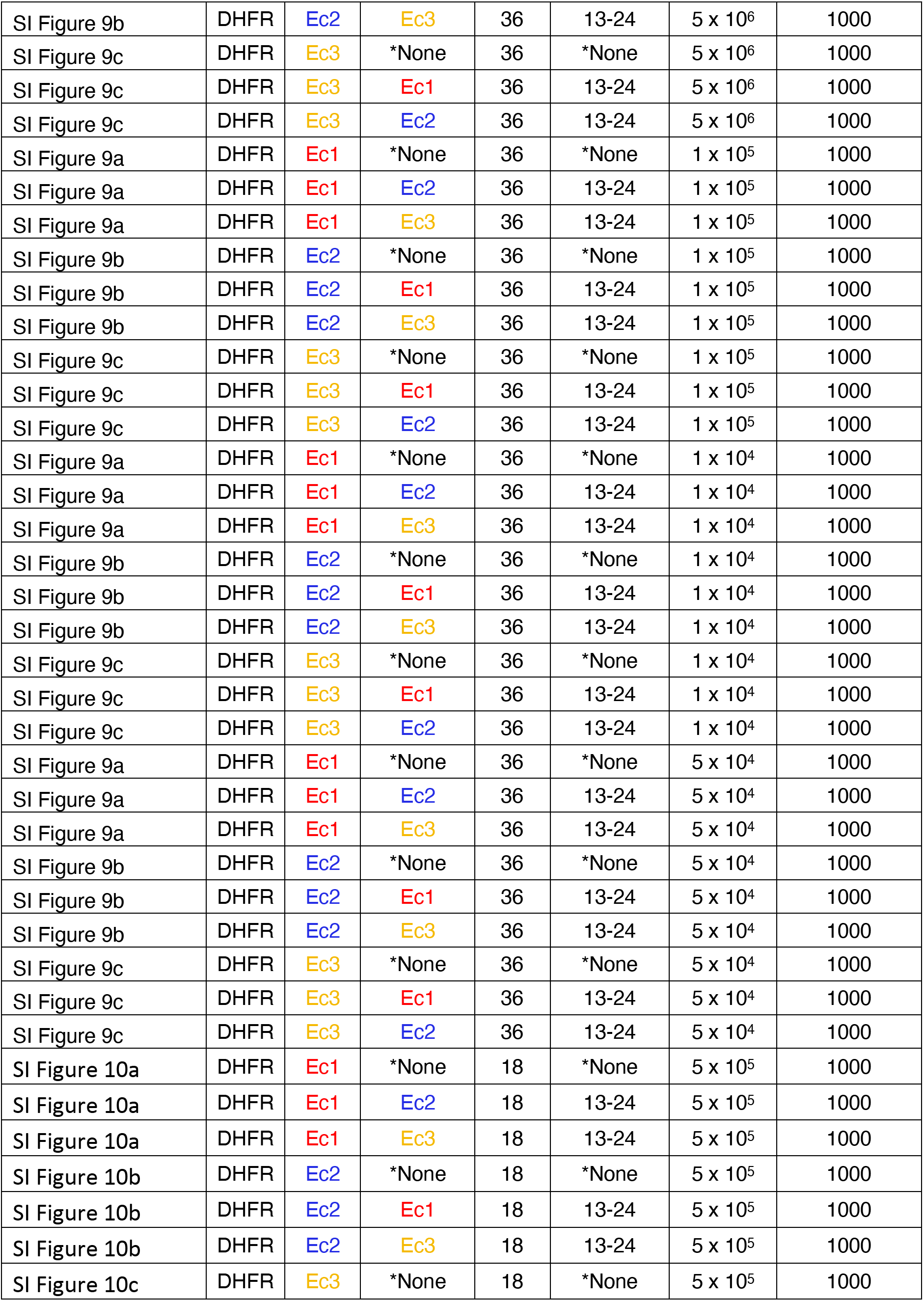
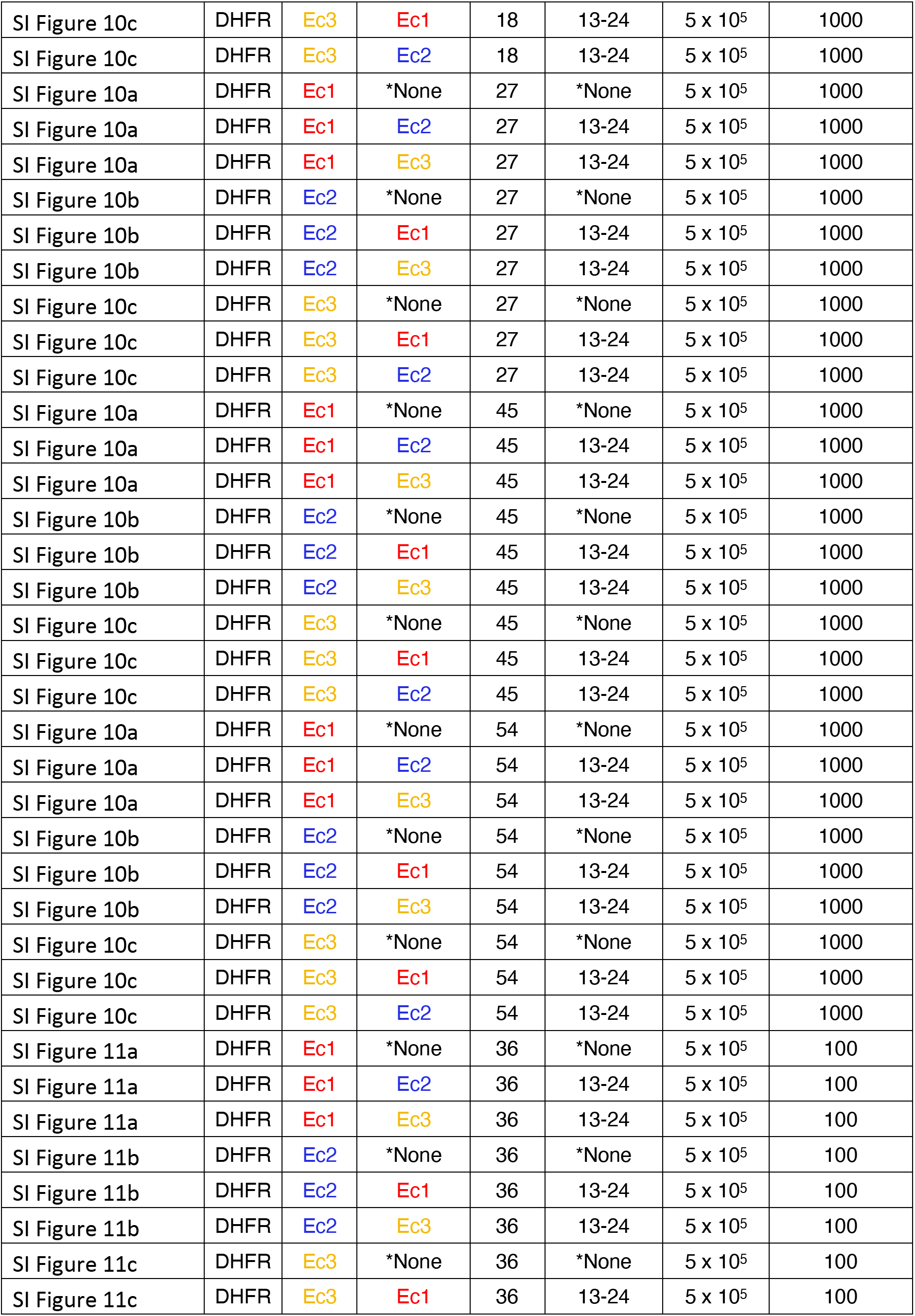
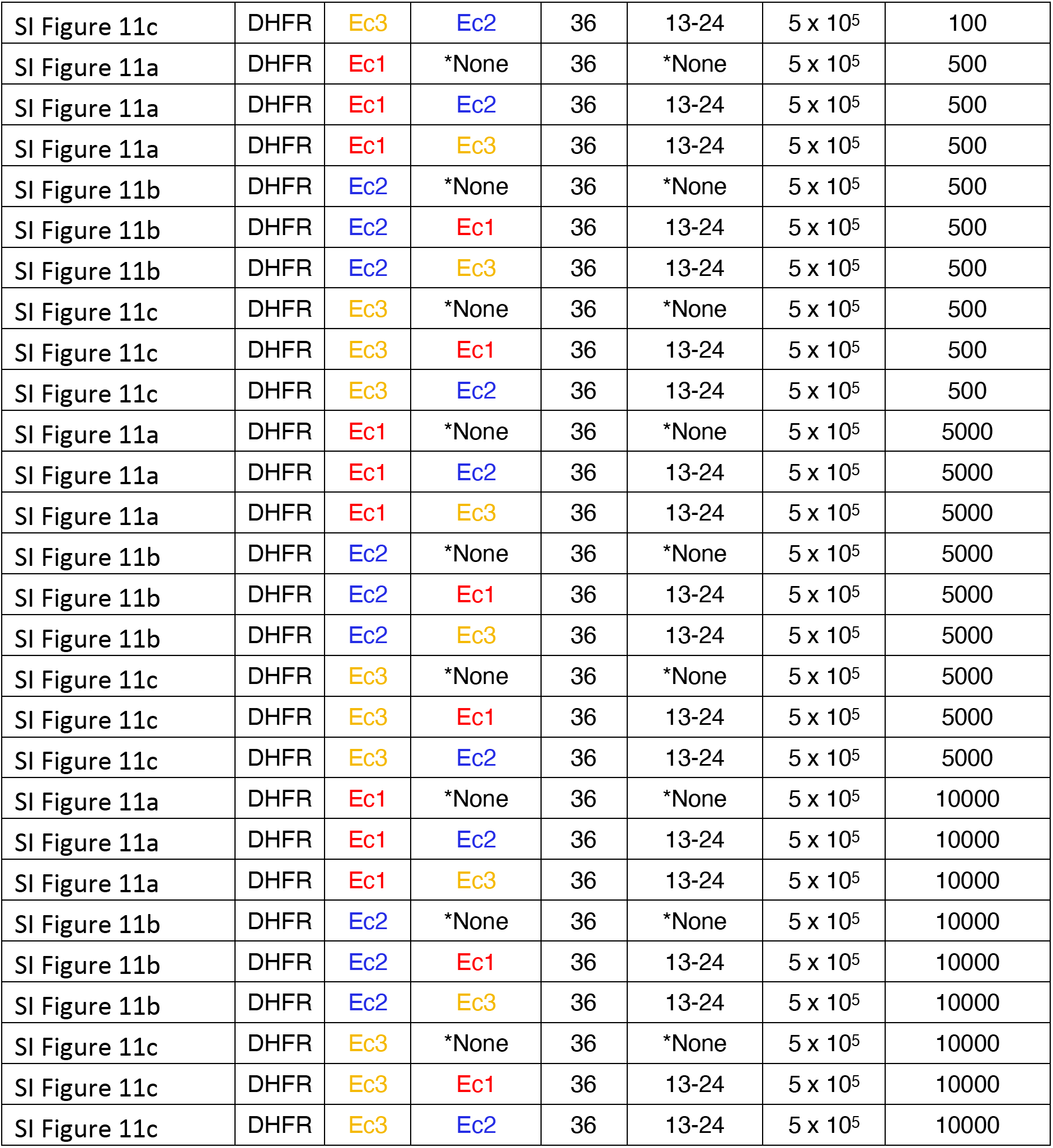
Specific datasets and parameters used in the evolutionary simulations. The same population size (1,000 individuals) was used in each treatment.

